# Neural Network-Derived Potts Models for Structure-Based Protein Design using Backbone Atomic Coordinates and Tertiary Motifs

**DOI:** 10.1101/2022.08.02.501736

**Authors:** Alex J. Li, Mindren Lu, Israel Desta, Vikram Sundar, Gevorg Grigoryan, Amy E. Keating

## Abstract

Designing novel proteins to perform desired functions, such as binding or catalysis, is a major goal in synthetic biology. A variety of computational approaches can aid in this task. An energy-based framework rooted in the sequence-structure statistics of tertiary motifs (TERMs) can be used for sequence design on pre-defined backbones. Neural network models that use backbone coordinate-derived features provide another way to design new proteins. In this work, we combine the two methods to make neural structure-based models more suitable for protein design. Specifically, we supplement backbone-coordinate features with TERM-derived data, as inputs, and we generate energy functions as outputs. We present two architectures that generate Potts models over the sequence space: TERMinator, which uses both TERM-based and coordinate-based information, and COORDinator, which uses only coordinate-based information. Using these two models, we demonstrate that TERMs can be utilized to improve native sequence recovery performance of neural models. Furthermore, we demonstrate that sequences designed by TERMinator are predicted to fold to their target structures by AlphaFold. Finally, we show that both TERMinator and COORDinator learn notions of energetics, and these methods can be fine-tuned on experimental data to improve predictions. Our results suggest that using TERM-based and coordinate-based features together may be beneficial for protein design and that structure-based neural models that produce Potts energy tables have utility for flexible applications in protein science.

**Code:** Code will be made publically available at https://github.com/alexjli/terminator_public

## 1 Introduction

A fundamental goal of computational protein design is to identify a protein sequence that can adopt a specified structure and exhibit a desired function^1^. Computational methods have guided the design of enzymes^2^ and small proteins that bind to the spike protein of SARS-CoV-2 and inhibit infection^3^. Traditionally, protein design has been dominated by physics-based models such as Rosetta^14^. However, the computational complexities of modeling molecular interactions at scale require judicious approximations of molecular physics. Accumulated error from these approximations results in inaccuracies, such that physics-based models typically must be coupled with multiple rounds of expensive experimental trial-and-error to meet design objectives ^1,3,5^.

Recently, deep learning methods have been applied to computational protein design and have achieved remarkable success on native sequence recovery benchmarks, where the task is to recapitulate the native sequence given the backbone coordinates of the associated structure. Most structure-based deep learning methods used for computational protein design represent proteins using static coordinates obtained from X-ray crystallography experiments^5,6,7^. Such representations typically respect the rotational and translational invariance of protein properties with respect to a coordinate basis and are generated by using the local environment to iteratively refine the features describing a residue or residue-residue interaction. A popular method of performing such computation is via graph neural networks (GNNs), which model proteins as a network of residues (nodes) and residue-residue interactions (edges)^5,6,8^. GNNs have achieved remarkable success on native sequence recovery tasks, significantly outperforming Rosetta on challenging test sets curated to be structurally distinct from the training data.

However, published models have several drawbacks. Neural network models can suffer from overfitting, a phenomenon in which the networks use a very large number of model parameters to memorize the training data rather than learning more general trends. As neural networks achieve increasingly better performance on native sequence recovery tasks, there is concern that models are becoming “backbone readers,” learning to read crystallographic patterns in published protein structures rather than extracting more general properties of protein structure that can be applied in new design problems.

Another drawback of current models arises from the form of output of the model. Natural language processing has spurred many advances in deep learning that have been subsequently applied in protein design. These frameworks generate sequences for input structures using a conditional per-residue learned probability distribution, which can be sampled from to generate new sequences either linearly (e.g., from N-terminus to C-terminus)^5,6^ or in a random order^8^. However, this type of sequential sequence generation is difficult to adapt to certain common protein design tasks. For example, such a distribution does not naturally allow for investigation of specific residue-residue interactions. It is costly and not intuitive to perform property-based optimization, such as designing proteins with net positive charge or high predicted solubility. More broadly, such models do not provide a picture of the sequence-function landscape of a protein structure, i.e., a map of how fitness changes as the sequence is varied, which is a property of great interest to protein scientists.

In light of these drawbacks, statistical models that capture sequence-structure relationships offer potential solutions. An approach focused on Tertiary Motifs (TERMs) has shown success quantifying relationships between sequence and structure^9^. TERMs are compact structural units that recur frequently in many unrelated proteins, and Mackenzie et al. showed that a large portion of protein structure space can be described using TERMs^10^. Information in TERMs can be used to define a statistical energy potential for a protein by searching for the closest matches to substructures in the protein across the PDB and quantifying the sequence statistics of the resulting matches. Such a potential function, in the form of a Potts model, can used in design, as implemented as a procedure known as dTERMen^9^. This procedure introduces a degree of backbone flexibility into design because there is permissiveness in the structural matching of TERMs. The overall approach and the dTERMen method in particular have proven valuable. Statistical models based on TERMs can detect incorrect regions in predicted structures^11^, predict mutational changes in protein stability^12^, and be applied directly to protein design^9,13^.

In light of the successes of dTERMen, we reasoned that features from this approach could be applied to build better GNN models for protein design. Specifically, introducing a structure featurization that doesn’t rely on a fixed set of backbone coordinates can potentially mitigate the problem of overfitting or coordinate memorization. Additionally, models that output an energy function over sequence variables can be used more flexibly and for a greater number of applications than models that directly output sequences. Potts models can efficiently describe an energy landscape over sequence space using a decomposition into single-residue and residue-pair contributions that is convenient and intuitive to structural biologists. Potts models can be used for tasks such as predicting mutational energies, optimizing just a subset of a structure, or sampling protein sequences under constraints^9,13^.

In this work, we designed two deep neural networks that generate an energy landscape over sequences on a particular structure: TERMinator, which takes both TERM data and backbone coordinate data as inputs, and COORDinator, which takes only backbone coordinate data. Comparisons of TERMinator and COORDinator show that TERM data are essential for the best performance. We additionally demonstrate that TERMinator designs are predicted by AlphaFold to fold to their target structures. Finally, we present evidence that both TERMinator and COORDinator learn notions of energetics, and we demonstrate that these models can be trained on experimental binding data to improve predictions of binding energies and protein stabilities. Our results suggest that TERMs provide a useful featurization of proteins for deep learning models and that outputting Potts models adapts GNNs for energy-based tasks.

## 2 Results

### 2.1 TERMs improve model performance on native sequence recovery tasks

#### Model Training and Datasets

TERMinator uses experimentally determined protein structures as training data to learn sequence-structure relationships. The model extends the Structured Transformer introduced by Ingraham et al. ^5,6^ and takes a structure as input to generates a Potts model over sequence variables as output; the loss function is the negative log of the composite pseudo-likelihood of the native sequence on a given structure, as described in the Materials and Methods. We trained TERMinator on two datasets: a previously-curated single-chain dataset split using CATH topologies^5^ (the “Ingraham Dataset”), and a new multi-chain dataset split using sequence redundancy (the “Multichain Dataset”). Details regarding model design, as well as the generation and composition of the two datasets, are discussed further in the Materials and Methods.

#### Ablation studies

For both datasets, we trained TERMinator and COORDinator, alongside several ablated versions, to understand how the different modules and input features affect performance on native sequence recovery (NSR). Native sequence recovery measures the percent residue identity between the original protein sequence and the sequence obtained by optimizing the structure-specific TERMinator-generated Potts model. Results are shown in Table 1 and detailed descriptions of each model ablation are in the Supplementary Information. Our best models improve upon the Structured GNN and also on the related GVP-GNN protein-design model of Jing et al., as demonstrated by performance on the Ingraham Dataset.^5,6^.

**TABLE 1.** Ablation Studies of TERMinator. Native sequence recovery is listed as mean ± standard deviation on median performance of triplicate train/test runs on the same data split, where triplicate data are available. dTERMen* is a handicapped version of dTERMen run without near-backbone TERMs. Underlined models indicate models released in this paper. Bold text indicates best performance among comparisons shown. TIC = TERM Information Condenser, GPME = GNN Potts Model Encoder: see Materials and Methods.

| Model version | Ingraham NSR | Multichain NSR |
| --- | --- | --- |
| <u>TERMinator</u>: TIC + GPME | **42.40 ± 0.02** | **46.16 ± 0.15** |
| Linearize TIC | 41.39 ± 0.16 | 44.56 ± 0.04 |
| Ablate TERM MPNN | 41.89 ± 0.11 | 45.42 ± 0.15 |
| Ablate Coordinate-based Features, Retain $k$-NN graph | 34.19 ± 0.05 | 37.89 ± 0.35 |
| Ablate GPME | 32.38 ± 1.65 | 36.62 ± 0.06 |
| <u>COORDinator</u>: GPME Alone (no TERM information) | 40.25 ± 0.17 | 43.46 ± 0.15 |
| dTERMen* | 24.32 | 26.61 |
| GVP-GNN<sup>6</sup> | 40.2 | - |
| Structured GNN<sup>5</sup> (as reported by Jing et al.<sup>6</sup>) | 37.3 | - |

Ablation studies indicate that although TERM-based features and coordinate-based features are largely redundant, neither serves as a full replacement for the other. COORDinator, which is trained purely on coordinate data and outputs a Potts energy function, achieves NSR on the Ingraham Dataset of 40.3%, outperforming the Structured GNN. Similarly, training on TERM data with no coordinate information (but with the benefit of the global *k*-NN graph, an inherently fuzzy feature) also achieves a respectable NSR of 34.2%. However, the full TERMinator model outperforms both of these models, as well as similar published models^5,6^, on NSR for the Ingraham Dataset. Similar trends in performance were observed for the Multichain Dataset.

When we remove coordinate data as inputs, TERMinator effectively acts as a better version of dTERMen^9^. Ablated TERMinator and dTERMen* (a handicapped version of dTERMen with no access to near-backbone TERMs) share access to essentially the same TERM input data (see the Supplementary Information for a more detailed discussion), but the learned model makes better use of this data, with an NSR of 34.2% vs. 24.3% on the Ingraham Dataset and 37.9% vs. 26.6% on the Multichain Dataset for TERMinator vs. dTERMen*.

### 2.2 TERMinator designs physically realistic sequences

#### TERMinator position-wise mutations are physically realistic

To better understand the performance of TERMinator, we examined what pointwise substitutions it makes when it does not predict the native residue in NSR tasks. This was done by examining the amino-acid confusion matrix, where the x-axis represents the predicted residue identity, and the *y*-axis represents the native residue identity. Representative matrices are shown in Figure 1. The strong diagonal in the matrix reflects the high overall native sequence recovery. Glycine (G), which is small and uniquely flexible, and proline (P), which lacks an amide proton and is conformationally constrained, are particularly well recovered. Interestingly, the substitutions the model makes are physically realistic. We see confusion within the EKR block, which contains charged amino acids glutamate (E), lysine (K), and arginine (R). The switch of charge polarity is potentially attributable to the model reversing the direction of salt bridges. Other blocks include the following: ST, with highly similar serine (S) and threonine (T) hydroxyl sidechains; VI, with sterically similar branched aliphatic valine (V) and isoleucine (I) sidechains; FWY, encompassing large hydrophobic aromatic residues phenylalanine (F), tryptophan (W), and tyrosine (Y); and DN, with isosteric aspartate and asparagine. These substitutions are highly plausible and suggest that TERMinator learns physicochemically realistic representations of proteins.

**FIGURE 1.**
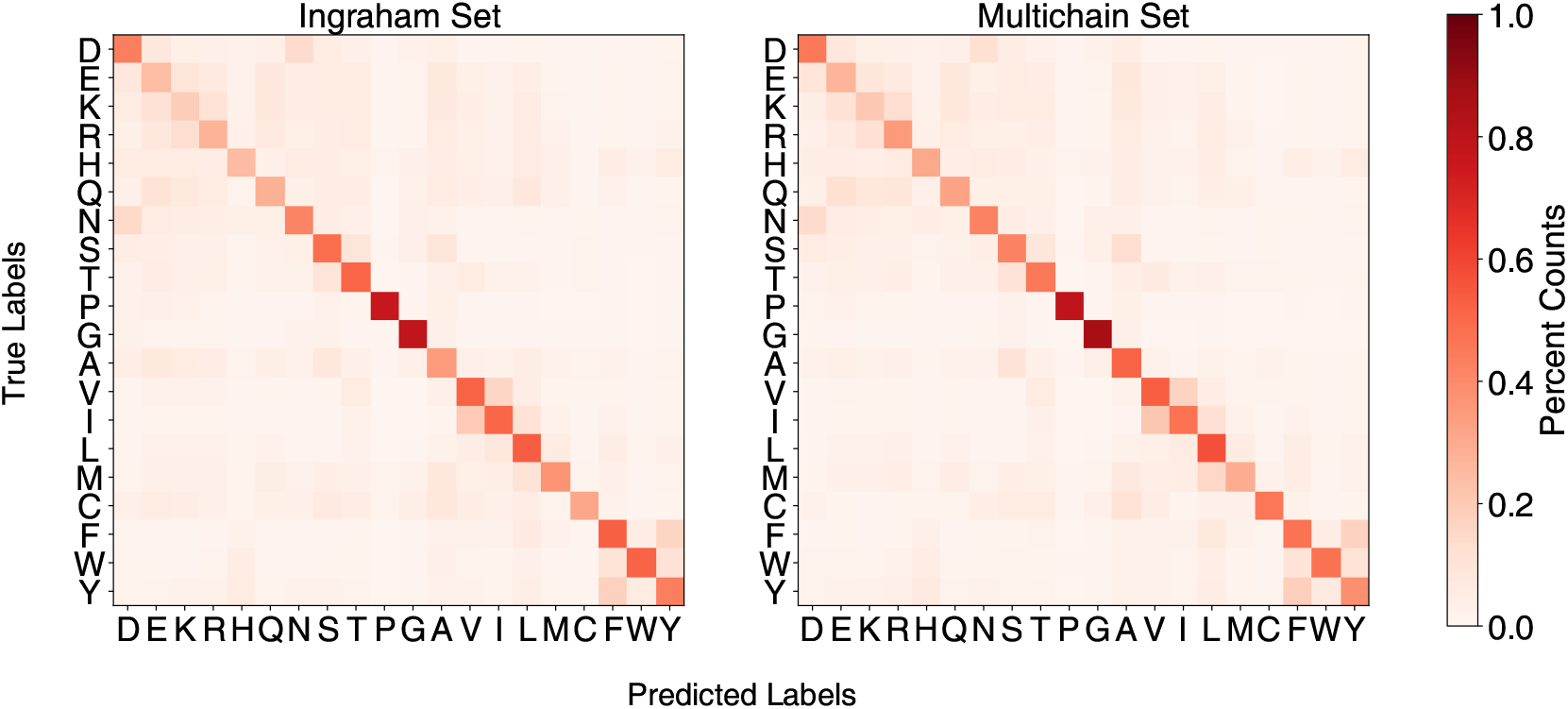
Confusion matrices comparing position-wise predicted residue identity (x axis) versus native residue identity (y axis). Values are normalized by the number of times the native residue occurs in the test set. Values report aggregate performance by the full TERMinator model across triplicate runs, on both the Ingraham and Multichain datasets.

#### TERMinator occasionally generates pathological, low-complexity designs

Although the confusion matrix reflects position-wise prediction trends, it does not capture sequence-wide properties. Upon examining TERMinator-designed sequences, we noticed that a non-trivial portion of designed sequences were dominated by one or a few residues. To detect such cases, we defined a complexity metric that counts the number of possible arrangements of residue labels in a designed sequence^14^ (see Materials and Methods for details). As shown in Supplemental Figure H1, the complexity distributions for native and TERMinator-designed sequences differed significantly: no native sequence in the Ingraham test set had a complexity value less than 1.67, whereas 198 of 1120 TERMinator designs had complexity score less than this value. Based on this, we classified designs with complexity < 1.67 as “low complexity.”

For low-complexity cases, we reran MCMC simulated annealing on the respective energy landscapes but added a complexity-based penalty during sequence design. After applying this penalty, the complexity of the designed sequences improved significantly, with no sequence having a complexity lower than 1.67 (Supplemental Figure H1). Among the redesigned sequences, we observed a 3.7% mean increase in NSR (Supplemental Figure H2). Given these results, we adapted our protocol to include re-design of any low-complexity sequences using the low-complexity penalty.

#### TERMinator-designed sequences are predicted to fold to the corresponding native structure

We evaluated whether TERMinator-designed sequences are predicted to fold to the target structure, a metric of designed-protein fold specificity. We used AlphaFold^15^ to predict the structure of TERMinator designs for the Ingraham test-set structures, all of which are single-chain structures. To evaluate performance, we used template modeling scores, or TM-scores^16,17^, which quantify the structural similarity of the predicted structure to the native structure. A TM-score of 0.7 corresponds to a ≥ 90% probability that the predicted structure is in the same fold family as the native structure. As expected, at high NSR, predicted structures matched the native fold very well. However, TERMinator also designed sequences with low NSR that were predicted to fold to the intended structure (Figure 2 A, 2 B).

**FIGURE 2.**
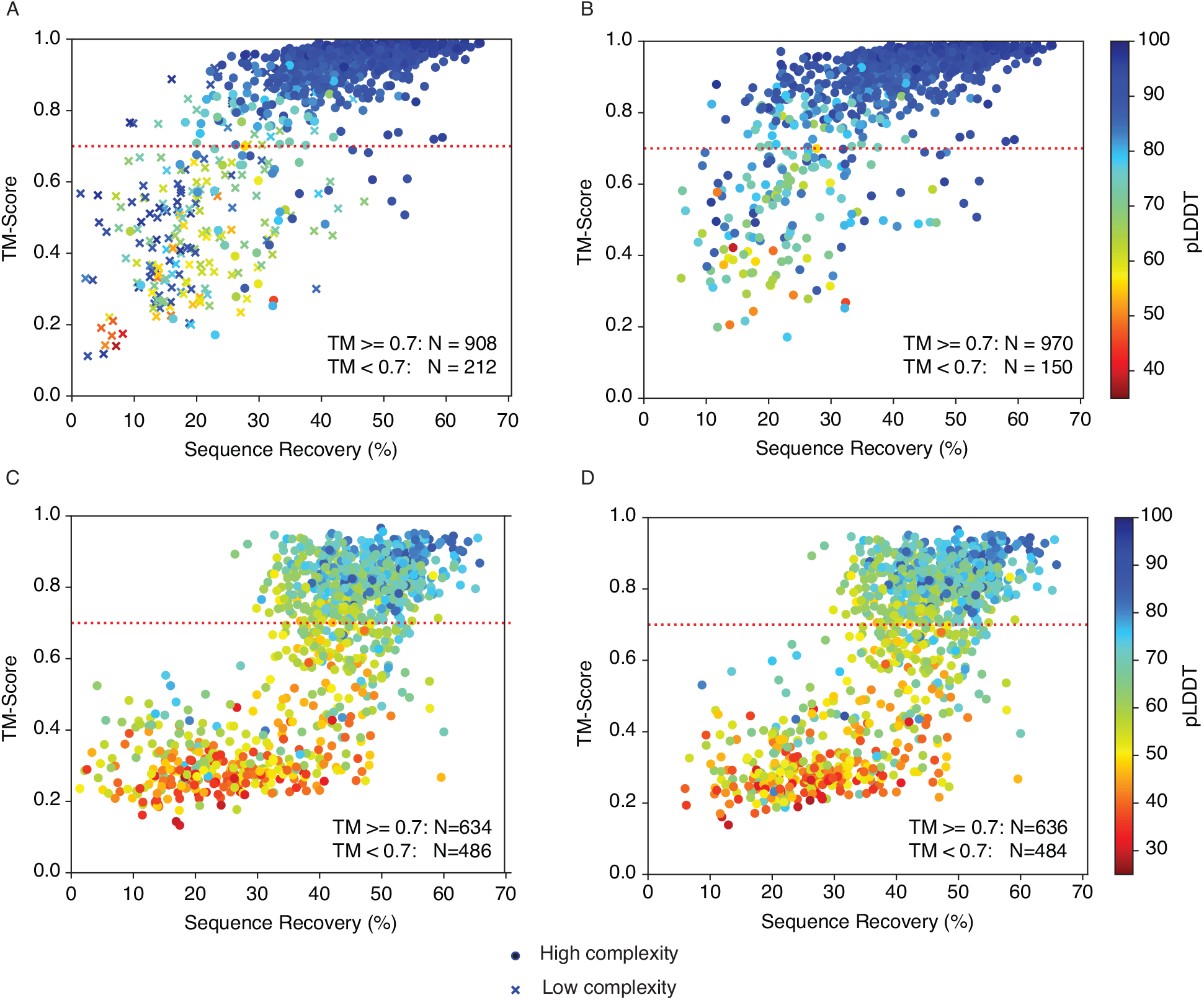
Fold-specificity analysis on Ingraham test-set cases. A) TM-scores for AlphaFold predictions for TERMinator-designed sequences versus native sequence recovery. Low-complexity sequences are indicated using crosses. B) Same plot as (A) after re-designing low-complexity sequences. C) Same plot as (A) but for random sequences with NSR values matching those in (A). D) Same as (C) but for random sequences with NSR values matching those in (B). In all panels, the AlphaFold predicted local distance difference test (pLDDT), which corresponds to confidence in prediction quality, is represented by the hue of the data point, as indicated in the heatmap key.

When 198 low-complexity designs were re-designed using the low-complexity penalty, we observed a 0.17 mean increase in TM-score, with 62 structures crossing the TM-score threshold of 0.7. (Figure 2 A versus 2 B). Notably, including the high-complexity cases, only 150 cases out of 1120 had TM-scores below 0.7, and only 70 had TM-scores below 0.5. Overall, the structures that AlphaFold predicted with high confidence generally matched the input structure fold; using the predicted local distance difference test (pLDDT) value as a metric for AlphaFold confidence, 924 of 975 predicted structures with pLDDT greater than 0.8 had a TM-score greater than 0.7.

To determine how much fold specificity TERMinator encodes beyond NSR alone, we created a control based on random sequences. We generated sequences with the same NSR values as the TERMinator designs, but with residues matching at random positions. Randomly generated sequences were folded using AlphaFold and TM-scores for the resulting structures were calculated with respect to the native chains (Figure 2 C, D). For random sequences with NSR greater than 50%, there is enough information that AlphaFold can often predict the corresponding fold, even though pLDDT is lower than for TERMinator-designed sequences. In contrast, for TERMinator-designed sequences, the high-NSR examples have a higher success rate, and many low-NSR sequences are predicted to adopt the target structure, as shown in Figure 2 B. These results, as well as the confusion matrices in Figure 1, suggest that the TERMinator potential can be used to provide physicochemically realistic sequences.

### 2.3 TERMinator and COORDinator can be used for analysis of protein energetics

TERMinator is not explicitly trained to model energetics. However, the output of the model takes the form of an energy function, and we investigated the correlation of TERMinator pseudo-energies with experimental observables including peptide binding affinities and protein resistance to proteolysis, which is a proxy for thermodynamic stability.

#### Binding affinities for Bcl-2 proteins

Following a benchmark performed with dTERMen by Frappier et al.^13^, we investigated whether Potts models generated by TERMinator and COORDinator could predict binding energies for Bcl-2 family protein-peptide complexes. Bcl-2 proteins are of particular interest due to their role in controlling apoptosis, a form of programmed cell death, and their overexpression in many cancers^18^. We compared our results to earlier work using other structure-based energy evaluations.

We used 13,084 affinity measurements across three anti-apoptotic Bcl-2 family proteins that bind to α-helical peptides of roughly 20 residues in length: Bcl-x_L_, Mcl-1, and Bfl-1^19^. Although the three proteins share the same fold, they have only 24% pairwise sequence similarity with one another^20^. We measured the Pearson correlation between TERMinator or COORdinator predicted energies and the experimental affinity measurements. The most relevant comparisons are shown in Figure 3, with the full results in Supplemental Table J1.

**FIGURE 3.**
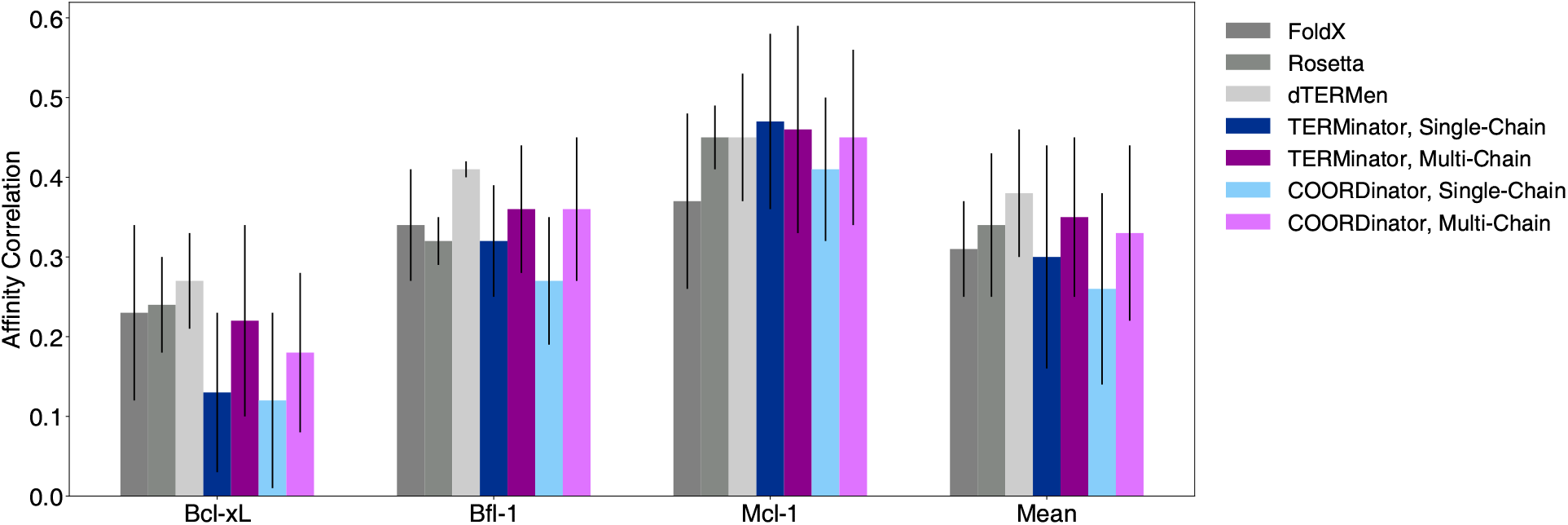
Structure-based prediction of binding energies for Bcl-2 protein complexes^19,13^. For multi-chain TERMinator and COORDinator, the Pearson correlation between the predicted and experimental binding energies was calculated and is reported as the mean and standard deviation over all structural templates and triplicate training runs, the variance arises from variable performance across structures, not replicates. The performance data for FoldX and Rosetta were taken from previous work^13^, and the performance of dTERMen was recomputed with the latest version of the model as of January 2022. Individual values can be found in Table J1 in the Supplementary Information.

The performance of TERMinator and COORDinator is on par with existing methods FoldX, Rosetta, and dTERMen that predict energies from structural data Frappier et al.^13^. Using TERMs in addition to backbone coordinates had only a small effect, which was not statistically significant for 2 of the 3 families. The models trained on the Multichain Dataset performed better than those trained on the Ingraham Dataset in all but one example (Mcl-1 predictions using TERMinator), which is unsurprising because the task involves predicting the stability of a two-protein complex (Supplemental Table J1). Using paired t-tests to compare models, and including data for all three proteins, we obtained *p* = 0.0011 between the multi-chain and single-chain forms of TERMinator and *p* = 4.68 × 10^−7^ between the two forms of COORDinator.

#### Stability of *de novo* mini-proteins

Next, we evaluated TERMinator and COORDinator on a dataset from Rocklin et al.^21^. In this work, the authors performed deep mutational scans for a set of *de novo* designed mini-proteins and three natural protein domains, Yap65, villin, and Pin1, using a high-throughput proteolysis assay that was shown to provide good estimates of protein stability^21^. We compared the results of single-chain and multi-chain TERMinator and COORDinator with the flexible-backbone version of the Structured Transformer of Ingraham et al.^5^, the GVP-GNN model of Jing et al.^6^, and the GVP-Transformer-AF2 model of Hsu et al^22^. These models were not all trained on the same data. The Structured Transformer and our models were trained on the Ingraham Dataset, which was derived from the CATH 4.2 annotations. The GVP models were trained using a similar dataset derived from CATH 4.3 annotations, and the GVP-Transformer-AF2 model used an additional 12 million structures predicted using AlphaFold^15^ as training data. We computed the Pearson correlation between TERMinator or COORdinator psuedo-energies and proteolysis-based stability values. We give the results for the 10 proteins benchmarked in previous work in Figure 4, and in Supplemental Table J2 we report performance on seven additional proteins in the Rocklin et al.^21^ dataset that were not included in previously published tests^22^.

**FIGURE 4.**
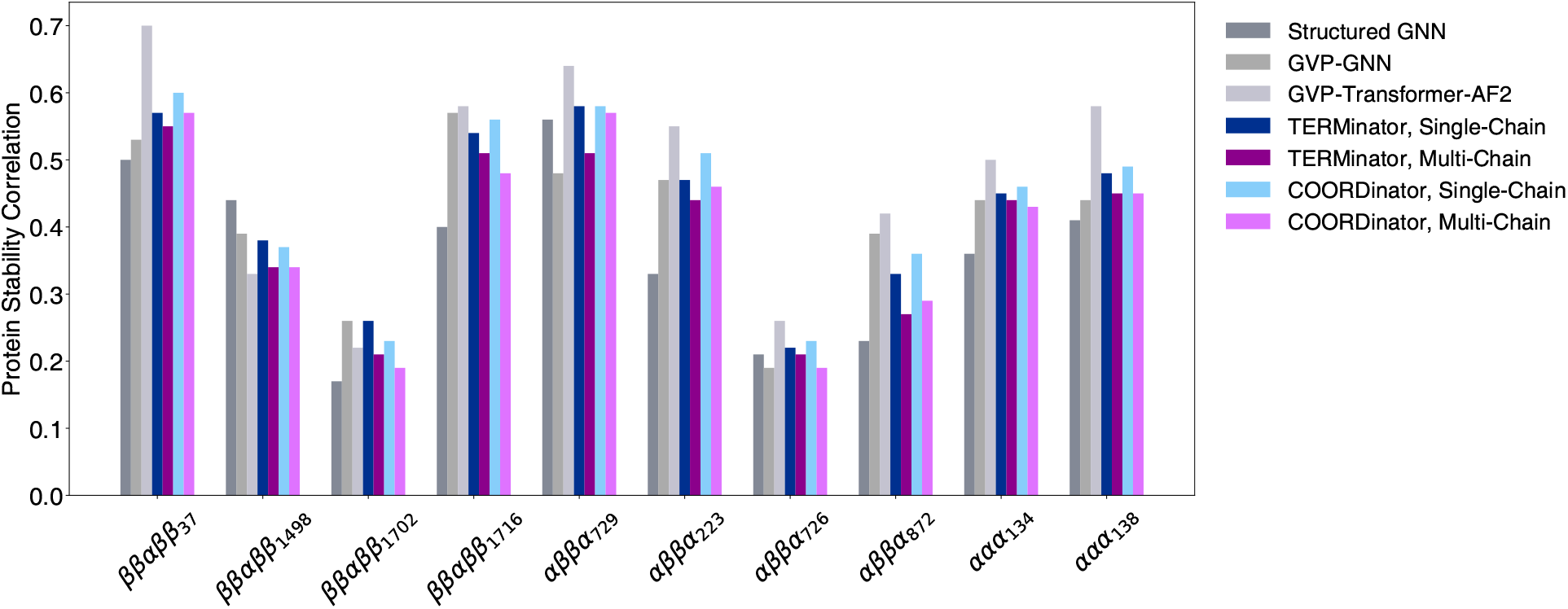
Structure-based prediction of mutational effects on designed protein domain stability^21^. The performance of TERMinator and COORDinator models is given as the average across triplicate training runs of each model, with standard deviations all ≤ 0.04 omitted for clarity. The performance of other models is reproduced from the literature. Individual values are available in the Supplementary Information in Table J2.

TERMinator and COORDinator both showed performance similar to other state-of-the-art models, aside from the GVP-Transformer-AF2 model, which performs the best but is also trained on 1000x more structures. Given the improvement in performance observed when the base GVP-GNN was augmented with additional training data using AlphaFold predictions, we expect that our models, which already outperform the GVP-GNN, may see a similar boost in performance after training data augmentation. Our single-chain models performed better than our multi-chain models, consistent with the test involving single-chain proteins. However, the addition of TERMs had little impact on the models’ predictive power for this mini-protein stability test.

#### TERMinator can be fine-tuned to improve energy-based protein analysis

Motivated by the results of our initial benchmarking, we tested whether incorporating experimental binding energy data in training could improve the performance of TERMinator and COORDinator on energy-based protein analysis tasks. We refined both models via a process known as fine-tuning, where a model that is already trained on a larger, more general dataset is updated in a re-training process that uses a smaller but more tailored dataset.

We used the Bcl-2 binding affinity data^19^ to fine-tune TERMinator and COORDinator. During fine-tuning, we trained the model to minimize a balance between the negative log composite pseudo-likelihood and the negative Pearson correlation between the experimentally measured binding affinities and predicted Potts model energies (see Materials and Methods).

Figure 3 shows that TERMinator and COORDinator perform best for Mcl-1 binding data, followed by Bfl-1 and Bcl-x_L_. For our fine-tuning test, we chose to use Mcl-1 binding data as the training set, as it had the best performance to begin with. We used Bcl-x_L_ data as the validation set; this allowed us to stop training before overfitting. Bfl-1 data served as our test set, on which we report results. We performed fine-tuning using both the single-chain and multi-chain models, and both TERMinator and COORDinator. The data in Table 2 show that fine-tuning is successful at improving affinity correlation performance on the held-out Bfl-1 family test set by an average of 0.19 points across all models.

**TABLE 2.** TERMinator and COORDinator performance before and after fine-tuning. Bfl-1 performance is reported as an average and standard deviation over all Bfl-1 complex templates across triplicate training runs. Performance on the Rocklin dataset^21^ is reported across triplicate runs over all 17 structures in the dataset. Native sequence recovery (NSR) is reported as the mean percentage test set recovery across median performance on triplicate runs on the Ingraham and Multichain datasets for the single-chain and multi-chain models, respectively. Per-fold performance on the Rocklin dataset can be found in Supplemental Table J3.

| Model version | Baseline |  |  | Fine-Tuned |  |  |
| --- | --- | --- | --- | --- | --- | --- |
| | Bfl-1 $r$ | Rocklin $r$ | Test NSR % | Bfl-1 $r$ | Rocklin $r$ | Test NSR % |
| TERMinator, Single-Chain | $0.32 \pm 0.07$ | $0.45 \pm 0.12$ | $42.40 \pm 0.02$ | $0.51 \pm 0.03$ | $0.53 \pm 0.11$ | $35.67 \pm 1.60$ |
| TERMinator, Multi-Chain | $0.36 \pm 0.08$ | $0.42 \pm 0.12$ | $46.16 \pm 0.15$ | $0.52 \pm 0.05$ | $0.51 \pm 0.11$ | $40.95 \pm 1.75$ |
| COORDinator, Single-Chain | $0.27 \pm 0.08$ | $0.46 \pm 0.13$ | $40.25 \pm 0.17$ | $0.51 \pm 0.03$ | $0.54 \pm 0.11$ | $33.52 \pm 0.99$ |
| COORDinator, Multi-Chain | $0.36 \pm 0.09$ | $0.42 \pm 0.14$ | $43.46 \pm 0.15$ | $0.51 \pm 0.04$ | $0.49 \pm 0.11$ | $38.28 \pm 0.42$ |

We also evaluated the models that were fine-tuned using Mcl-1 and Bcl-x_L_ protein binding data on the Rocklin dataset. This represents a stringent test of generalizability, because the Rocklin dataset contains *de novo* designed structures that have no relation to the Bcl-2 family. In Table 2, we show that fine-tuning improved performance on this test by an average of 0.08 points across all models. This improvement suggests that we successfully finetuned TERMinator and COORDinator in a way that generates Potts models that are more physically realistic and potentially broadly useful. For the single-chain models, numerical improvement results for each individual fold are in Supplementary Table J3.

We evaluated the fine-tuned single-chain and multi-chain models on the original Ingraham and Multichain test sets, respectively, and report NSR performance in Table 2. The NSR values for the multi-chain models each dropped about 5%, while the NSR on single-chain models fell about 7-9%. Thus, although performance on energy benchmarks improved after fine-tuning, it came at a cost to NSR. Note that the low-complexity cases obtained from fine-tuned versions of all single-chain and multi-chain models were re-designed with the same protocol discussed in Section 4.4.

### 2.4 Investigation of TERMinator Potts models reveals disconnect between sequence-based and energetics-based metrics

As discussed above, we noticed that TERMinator and COORDinator Potts models sometimes assigned low energies to low-complexity sequences (e.g. long repeating stretches of the same residue label). Although this did not occur for the majority of structures, and we were able to avoid this behavior by imposing a penalty on low-complexity sequences during design, we were curious about why the learned Potts models favored these clearly incorrect solutions.

Given dTERMen does not exhibit the same preference for low-complexity sequences, we compared the single-residue and residue-pair terms in the dTERMen vs. TERMinator and COORDinator Potts models. The scatterplots in Supplemental Figure K1 reveal two clusters corresponding to the self-energy parameters (orange) and the pair-energy parameters (blue). Notably, TERMinator puts more weight into the pair-energy parameters, whereas dTERMen puts more weight into the self-energy parameters.

We speculated that the relatively larger pair-energy weights in TERMinator may allow certain Potts models to have pathological low-energy, low-complexity minima that arise from spurious second-order effects. Hence, we sought to encourage TERMinator to place more weight on self energies in training. Due to the much greater number of pair-energy parameters compared to self-energy parameters, we decided to train TERMinator with a penalty on the vector norm of all learned parameters (see Materials and Methods), as a Potts model with larger self energies but smaller pair energies would have a smaller norm than a Potts model with the opposite characteristics. We then performed design without any low-complexity penalty. Notably, using the re-trained model, not only did the complexity of the sequences designed for test-set structures increase (Supplemental Figure H3), but the median NSR also increased by as much as 2.5% (Table 3).

**TABLE 3.** TERMinator and COORDinator performance with and without the Potts model parameter norm penalty. Native sequence recovery is listed as mean ± standard deviation over median performance for triplicate train/test runs on the same data split. For the Bcl-2 results, we report the performance averaged across all families (Mcl-1, Bfl-1, Bcl-x_L_) using the multi-chain version of each model. For the Rocklin results, we report the performance averaged across all 17 structures in the dataset using the single-chain version of each model.

| Model version | Ingraham NSR | Multichain NSR | Bcl-2 $r$ | Rocklin $r$ |
| --- | --- | --- | --- | --- |
| TERMinator, no norm penalty | 42.40 ± 0.02 | 46.16 ± 0.15 | **0.35 ± 0.10** | 0.45 ± 0.12 |
| TERMinator, with norm penalty | **44.52 ± 0.21** | **48.64 ± 0.02** | 0.33 ± 0.15 | 0.41 ± 0.11 |
| COORDinator, no norm penalty | 40.25 ± 0.17 | 43.46 ± 0.15 | 0.33 ± 0.11 | **0.46 ± 0.13** |
| COORDinator, with norm penalty | 42.24 ± 0.18 | 46.02 ± 0.12 | 0.32 ± 0.14 | 0.40 ± 0.11 |

We expected that the re-balanced Potts model would also improve TERMinator’s performance on the Bcl-2 and Rocklin binding and stability benchmarks. However, this was not the case. As shown in Table 3, models trained using the penalty did not give improvements on either task.

## 3 Discussion

### Ingraham vs Multichain Dataset

Our study builds on the pioneering formulation of protein design using the Structured Transformer, as proposed by Ingraham et al^5^. We have explored several modifications and extensions of this work, including training the model to learn a Potts model of protein energy as a function of sequence variables and incorporating a “fuzzier” representation of sequence-structure relationships using TERMs. Another change was our use of different training data for modeling protein complexes. Other recent work by Dauparas et al. ^8^ has also expanded the Structured Transformer using different approaches, e.g., investigating different backbone featurizations and decoding schemes.

To train TERMinator and COORDinator, we used two datasets. The Ingraham Dataset was generated by splitting all proteins in the non-redundant CATH 4.2 dataset into their individual chains, and then partitioning the chains into three sets such that no CATH fold was present in more than one set. This process included many protein chains without the binding partners (other protein chains) that are present in experimentally studied complexes. Because a binding partner may affect the protein conformation or structural environment of residues in a single-chain training example, such cases of missing contextual information may reinforce non-physical sequence-structure relationships, thereby compromising the ability of a model to describe binding interfaces correctly.

We generated our Multichain Dataset by splitting a non-redundant set of the PDB into three sets such that no individual chain in the test set shared more than 50% sequence similarity with any chain in the training or validation datasets. Although this led to a less rigorous structural split amongst the three sets than in the Ingraham Dataset, because training and test examples may have the same CATH fold, this strategy allowed the model to learn about all protein chains in the context of their binding partners.

TERMinator and COORDinator performed better on the Bcl-2 benchmark when trained on the multichain dataset. This is expected because the Bcl-2 dataset involves a binding interface between a peptide and a target protein. In contrast, all structures in the Rocklin benchmark are single-chain structures, and TERMinator and COORDinator performed better on this test when trained on single-chain data, possibly benefiting from the stricter training split and improved generalization.

### TERMinator vs. COORDinator

TERMinator and COORDinator, while built from the same modules, differ in that TERMinator has access to both TERM and coordinate data whereas COORDinator uses coordinate data alone. Notably, TERMinator achieves higher NSR than COORDinator (Table 1,3), which suggests that on tasks where recapitulating the native sequence is a good metric, TERMs provide non-redundant information.

However, mining TERMs is relatively time-expensive, at roughly 4 min per residue. While our studies indicate that TERMinator outperforms COORDinator on NSR, TERMs do not consistently improve energetics on either the Bcl-2 or Rocklin benchmarks. Therefore, in applications that require the evaluation of many structures, mining TERMs may not be worth the additional overhead cost. In such cases, COORDinator may be of greater use, as it performs well on the tested benchmarks without requiring TERMs. COORDinator has the potential to enable more facile high-throughput *in silico* energetics analyses: energy tables can be generated in seconds, with an average inference time of 73 *μ*s per residue using 1 NVIDIA V100 GPU and 465 *μ*s per residue on CPU.

### Benefits of Potts models for representing energy landscapes

Given that TERMinator is an energy-based model, it can be applied to any question that can be formulated in an energy-based framework, e.g. questions related to binding affinity or mutational effects. Additionally, the energy function is decomposable into component interactions, enabling investigation of and optimization over specific residues or reside-residue interactions. The Potts model form also allows for various other tasks such as constrained optimization, as done with dTERMen and other Potts models^13,19,23^. This opens up many possibilities for design, especially if we consider the additional potential for model improvement via fine-tuning and additional training data.

### Reflecting on metrics in protein design

During our investigations of model performance, we noticed several phenomena that raise interesting questions regarding Potts model design and the suitability of using NSR as a proxy metric for protein design.

First, although the median NSR of TERMinator and COORDinator is high, compared to other methods, the model occasionally produces low-information sequences with very low native sequence identity. Notably, the structures that give rise to low-complexity sequences are, themselves, often of low structural complexity. For example, the Ingraham test set contains single chains from helical bundles and coiled-coil oligomers that are relatively featureless alpha helices. In our NSR tests, sequences for these helices were designed without the appropriate oligomeric context. While such structures’ native sequences are not low-information, the decontextualized input helps explain why TERMinator-generated sequences have this quality.

To investigate this phenomenon in more detail, we examined the distribution of energetic parameters of TERMinator relative to statistically-derived dTERMen. After observing that the TERMinator Potts model distributes energetic parameters differently from dTERMen Potts models, we attempted to prevent pathological cases by implementing a penalty on the vector norm across all energetic parameters. Although this increased NSR across the board, no performance increase was seen on energetics-based tasks. Given that protein stability and binding energy are much more important than NSR for most protein design tasks, this casts doubt on the suitability of NSR as a metric when performing method development.

Another disconnect between NSR and desired model properties was observed when we folded TERMinator-designed sequences using AlphaFold ^15^. A number of designed sequences had relatively low NSR values but high TM-scores, reinforcing the point that sequences with low similarity to the native sequence can still fold to the desired structure. Given that variation on the native sequence may be required for many applications (for example, increasing or decreasing the immunogenicity of a protein, or increasing its solubility while maintaining its structure), selecting for protein design models based on high NSR may limit design flexibility. A similar criticism applies to selecting models based on low model perplexity.

### Promise of improving models with additional data

TERMinator and COORDinator are promising models because of the capacity to improve predictions given additional data. One avenue for improving the performance of protein design models is to allow them to learn directly from experimental data. There are a limited number of experimental datasets large enough for machine learning purposes, but our proof-of-concept shows that fine-tuning TERMinator on the Bcl-2 affinity benchmark improved performance not only for Bcl-2 binding predictions but also for the unrelated Rocklin proteolysis stability benchmark. This indicates the capacity for model generalization and the potential of making larger improvements by incorporating other data.

Additionally, with the advent of AlphaFold^15^, the use of predicted structures as training data provides a promising way to boost the performance of protein design models. Scaling the size of the TERMinator model and expanding the training data using predicted structures (as done in Hsu et al.^22^) could lead to a significant performance enhancements and greater generalizability.

## 4 Materials and Methods

### 4.1 Datasets

We trained TERMinator on two datasets: a published single-chain dataset^5^, and a new multi-chain dataset. For every protein, we described the structure using a set of Tertiary Motifs (TERMs), which are small, local-in-space fragments of proteins that recur frequently across unrelated proteins^10^. The selected TERMs consisted of sequence-local singleton TERMs (up to 6 contiguous residues) and sequence non-local pair TERMs (usually 2 interacting 3-residue segments). For each TERM, we performed substructure lookup^9^ against a non-redundant database generated from the Protein Data Bank (PDB) on Jan. 22, 2019. Self-matches with the protein itself were discarded. For each match, we recorded the sequence along with several geometric features of the backbone. For more details about the recorded TERM data, see the Supplementary Information.

Each dataset was split into training, validation, and test subsets, as described below. The model was allowed to learn from the training data. The validation data were used to determine when to terminate training so as to avoid over-fitting. Model performance was evaluated using the test dataset.

#### Multichain Dataset

We constructed the multichain dataset in the following manner. First, the PDB on Jan. 22, 2019 was filtered for X-ray structures primarily containing protein components, such that no chain in any given structure contained more than 500 residues. Then, membrane proteins were removed, and structures were filtered for resolution better than 2.6 Å. This resulted in 115583 proteins. This set of proteins was made non-redundant using usearch^24^ by clustering structures based on sequence similarity and then sampling one representative structure from each cluster. This resulted in 14747 protein structures. This set was then randomly split 90/10 into a training subset and a validation subset, both of which were filtered to remove any structures that contained fewer than 30 residues. This resulted in 13083 training-set structures and 1455 validation-set structures. The test data were constructed from all structures in the PDB deposited between Jan. 22, 2019 and May. 12, 2021. We computed sequence redundancy across all chains in the newly deposited structures by running BLAST+^25^ with an E-value criterion of 10^−6^ against the training set, validation set, and all structures used in the TERM match lookup database. We then discarded structures for which at least one chain within the structure had >50% computed sequence redundancy. The filtered dataset was then made non-redundant by using usearch and sampling one representative structure from each cluster; all structures that contained fewer than 30 residues were removed. This resulted in 1105 test-set structures.

### 4.2 Architecture

Our architecture is based on the Structured Transformer from Ingraham et al. ^5^, which we adapted to consider new inputs and produce Potts models as outputs. At a high level, TERMinator generates a Potts model by iteratively refining a graph-based representation of a protein. The network, shown in Figure 5, can be broken into two sections. The first section, the TERM Information Condenser, learns local structure via graph updates on small, local TERM graphs. The second section, the GNN Potts Model Encoder, learns global structure via graph updates over nearest neighbors across the entire chain. Using this global structure graph representation, the model outputs a Potts model over positional residue labels. COORDinator operates in a similar fashion, but consists only of the GNN Potts Model encoder. More information about the network architecture can be found in the next section and in the Supplementary Information.

**FIGURE 5.**
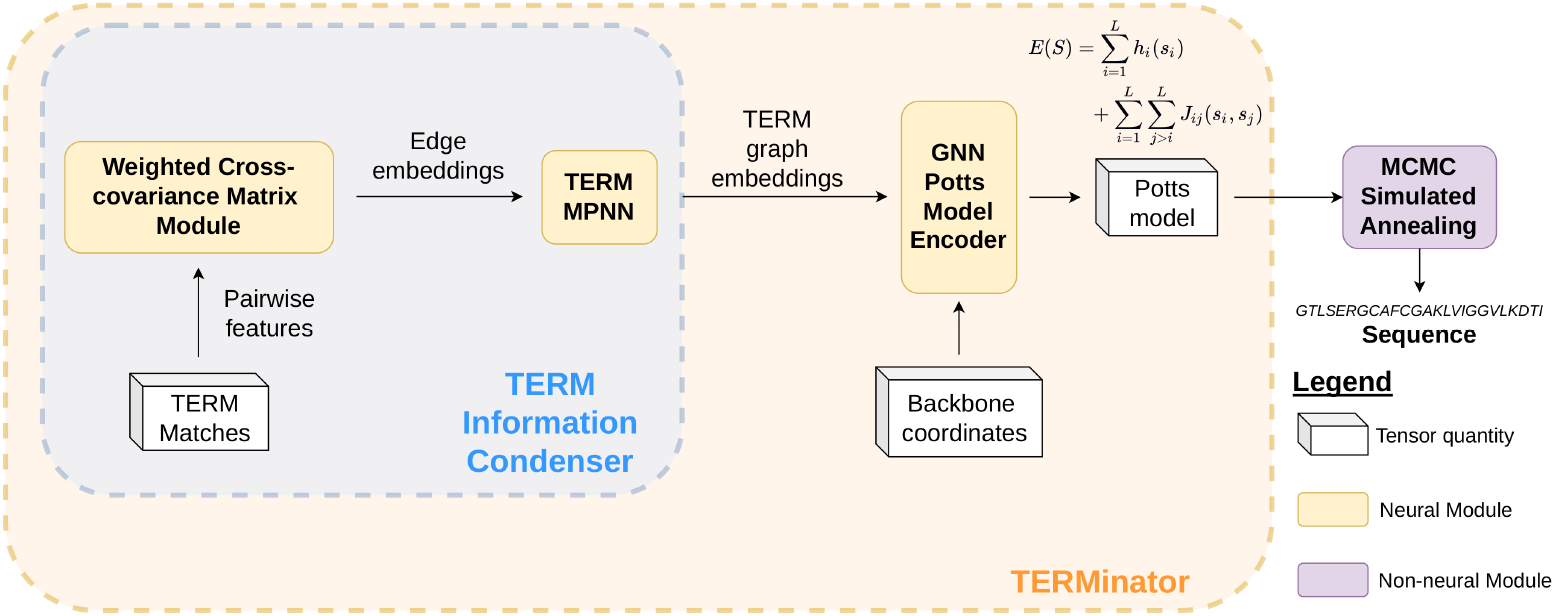
Model Architecture. The TERM Information Condenser extracts information from structural matches to TERMs in the target protein to construct node and edge embeddings. The GNN Potts Model Encoder takes in TERM data and coordinate features and outputs a Potts model over positional amino acid labels (see Supplementary Information: Potts Model for functional form). We use MCMC simulated annealing to generate optimal sequences given the Potts model.

#### TERM Information Condenser

TERMinator decomposes a target structure into TERMs and mines structural matches to these TERMs (as described above in Section 4.1). We represent the TERMs as bidirectional fully-connected graphs that include self-edges. Per TERM, we take the top 50 matches in the PDB with lowest RMSD. Each residue in each TERM match is converted to a set of geometric features describing backbone geometry and residue accessibility along with a one-hot encoding of the match residue identity. Node embeddings are initially set as an empty zero vector of proper dimensionality. Edge embeddings, representing residue interactions within a TERM, are computed using cross-covariance features between residues in a TERM. Self-edges are also represented in this manner.

We feed these preliminary TERM graph embeddings through a neural module called the TERM MPNN, which creates a featurized graph representing each TERM in a protein. Via a series of learnable graph updates, a collection of residue embeddings and edge embeddings per TERM is generated. We then merge all TERM graphs to form a full-structure graph, taking advantage of the fact that all residues and residue interactions in the structure are covered by at least one TERM. For nodes and edges that are covered by multiple TERMs, the representation for that node or edge is generated by averaging the representations across all covering TERMs.

#### GNN Potts Model Encoder

The GNN Potts Model Encoder combines the structure embedding produced by the TERM Information Condenser with target protein backbone coordinate features to produce a Potts model over sequence space. TERMinator uses the coordinate-based structure embedding presented in Ingraham et al.^5^, and this is fused, via concatenation of features, with the TERM-based structure embedding from the TERM Information Condenser. Concatenation of the coordinate-based and TERM-based embeddings occurs if the node or edge exists in the global *k*-nearest neighbors graph of residues; otherwise, a zero-vector of equivalent dimensionality is concatenated to the coordinate-based embedding. COORDinator uses only the Ingraham coordinate-based structure embedding.

The initial full-chain representation is processed by the GNN Potts Model Encoder (see the Supplementary Information for further details). The GNN Potts Model Encoder is a neural module that is identical to the TERM MPNN in architecture, but which operates on the global *k* nearest-neighbors (*k*-NN) graph, including self-edges. We produce a Potts model from the output of this network by projecting each edge embedding into a matrix of residue-pair interaction energies, averaging any duplicate bidirectional interaction energies (e.g. interaction *i* → *j* and interaction *j* → *i*). Self-energies are defined as the diagonal of this matrix (derived from the self-edge), whereas pair-energies are defined by the entire matrix.

### 4.3 Training

#### Loss Functions

The primary loss function used was the negative log *composite pseudo-likelihood* of the native sequence on a given structure, averaged across residue pairs. Composite pseudo-likelihood is the probability that any pair of interacting residues has the same identity as that pair of residues in the target sequence, given the remainder of the target sequence. Composite pseudo-likelihood can be defined using the energies described by the Potts model as follows. As stated in the Supplementary Information section “Potts Model,” a Potts model is defined by two functions: the self-energy function *E_s_*(*R_i_* = *m*) evaluates the energy of residue *i* with identity *m*, and the pair-energy function *E_p_*(*R_i_* = *m*, *R_j_* = *n*) evaluates the energy of residue *i* with identity *m* interacting with residue *j* with identity *n*. From the Potts model, we computed the contextual pairwise energy *E_cp_* for a sequence with residue *i* having identity *m* and residue *j* having identity *n*, provided all other residues *R_u_* with identity *r_u_*, as:

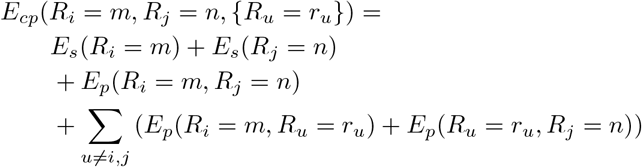

From this energy, we compute the composite pseudo-likelihood *p*(*R_i_* = m, *R_j_* = *n*, {*R_u_* = *r_u_*}):

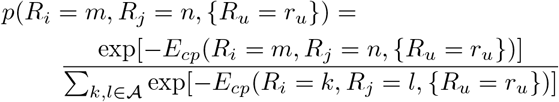

where 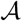 represents the set of all residue identities.

During general model training, the loss function was the negative log composite pseudo-likelihood averaged across residue pairs.

The Potts model norm penalty takes the form

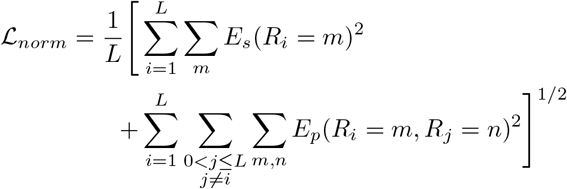

This is equal to the normalized L2 norm of all energies in a given Potts model. To train models with this norm penalty as discussed in Section 2.4, we used a weighted sum of the negative log composite pseudo-likelihood and 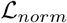 as the loss function. Using a logarithmic hyperparameter search, we empirically found that weighting these two terms equally led to the best NSR performance on the validation set; this choice was then used for training all evaluated models.

The fine-tuning objective used the negative Pearson correlation. Consider *N* sequences in a binding affinity dataset. For the *i*th data point, let the experimental binding value be *e_i_* and the predicted binding value be *p_i_*. We sought to minimize the negated Pearson correlation, or equivalently maximize the Pearson correlation, resulting in

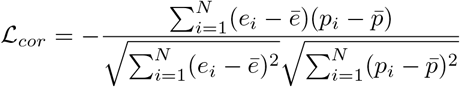

To fine-tune our models, we used a weighted sum of the negative log composite pseudo-likelihood and 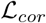 as the new loss function. Using a logarithmic hyperparameter search, we found that placing 100 times more weight on 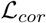 gave the best compromise between substantial improvements on the Bcl-2 validation set (Bcl-x_L_) and modest loss of NSR performance on the Ingraham and Multichain datasets. This choice was used for all fine-tuning.

#### Training Scheme

We trained each model for 100 epochs with early validation stopping, using the Noam optimizer as reported in Vaswani et al. ^26^ based on a model dimensionality of 128. We used a semi-shuffle batching method which operates in the following fashion: all proteins are first sorted according to their number of TERM residues, then partitioned into partitions of size 500. Within each partition, the order of the proteins is fully shuffled. Then, variable-size batches are constructed starting with the protein with the fewest number of TERM residues by incrementally adding proteins to a batch as long as the batch has less than 55000 TERM residues, after which a new batch is started and generated in similar fashion. For COORDinator, we perform the same process but add to the batch while it has less than 6000 residues. After all batches are constructed, the order of batches is shuffled fully before training begins. This process was repeated after every epoch.

The intuition behind this training scheme is to achieve a balance between full shuffling and compute efficiency. We sought to sort proteins of similar length together in order to prevent wasted computation on padding values. However, training on fixed mini-batches may lead to poor training convergence, so we also wanted mini-batches to have variability between training epochs. Thus, we opted to shuffle and batch proteins with other similar-length proteins.

### 4.4 Sequence design

To design sequences, given the Potts model for a structure, we used Markov Chain Monte Carlo (MCMC) processes for global optimization. For each Potts model, we sampled 100 sequences via MCMC-simulated annealing, running 10^6^ cycles per run while reducing the temperature from *kT* = 1 to *kT* = 0.1. The lowest energy sequence was then taken as the designed sequence.

If a designed sequence was designated as low-complexity (as described in 2.2, see Eqn. 4 of Wootton and Federhen^14^), we re-designed it using a low-complexity penalty. Specifically, we computed the number of unique arrangements of the amino-acid labels in sequence *S* as:

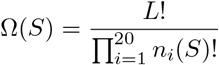

where *n_i_* is the number of times amino acid *i* occurs in *S*. We then defined the complexity of *S* as:

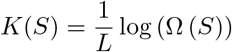

and if it was below 1.67, we labeled the sequence as low-complexity and added a penalty term to the Potts energy *E_Potts_* to generate a composite score:

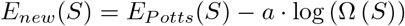

which was used in MCMC simulated annealing. We empirically found *a* =1 to work well. We ran redesign MCMC by generating 10 sequences, running 2 × 10^6^ cycles for each while reducing the temperature from *kT* = 10 to *kT* = 0.1. We chose the lowest-energy sequence (according to *E_new_*) as the final design.

### 4.5 AlphaFold folding

The sequences designed by TERMinator for the Ingraham test-set structures were folded with AlphaFold^15^ using the full-database parameter and default settings in monomer mode. Folding failed for a single case (Chain A of 3CNX) due to a failure in featurizing potential templates. This case was re-run on ColabFold^27^, which utilized a protein database with 70% sequence identity filter for searching for templates and the MMseq2 ^28^ sequence search tool. After folding, the TM-Score^16,17^ of the model with the highest pLDDT was calculated against the native structure.

Randomized sequences were generated by retaining the native residue identity at randomly selected positions and mutating the remaining positions to a randomly selected non-native residue. More specifically, 1) the number of residues to set as native was determined based on the NSR of the TERMinator design (rounded up to the nearest integer), 2) the positions to retain as native were selected randomly, in the new semi-random sequence, and 3) the rest of the residues were assigned at random from the 19 residues not equal to the native residue. This resulted in randomized sequences with native-sequence identities that matched those of the corresponding TERMinator designed sequences.

### 4.6 Energetics and Fine-Tuning

#### Bcl-2 Benchmark

As was done with dTERMen in Frappier et al.^13^, we used a dataset of 4488, 4648, and 3948 affinity measurements for BH3 peptides binding to Bcl-x_L_, Mcl-1, and Bfl-1, respectively^19^. Multiple structures are available for each protein in complex with different helical peptides. Following Frappier et al.^13^, we used 15, 25, and 6 structures for Bcl-x_L_, Mcl-1, and Bfl-1, respectively, as inputs to generate Potts models using TERMinator or COORDinator. To make binding energy predictions, we summed the Potts model self-and pair-energy contributions for all peptide positions. Per template, we computed the Pearson correlation between these values and the experimental affinity measurements. We then aggregated results per family by computing the mean and standard deviation across templates within a family.

#### Rocklin Benchmark

We used a dataset from Rocklin et al. ^21^ that was generated by performing deep mutational scans and using a proteolysis assay to quantify fold stability in high throughput. The authors studied a set of *de novo* designed mini-proteins and three natural protein domains, Yap65, villin and Pin1, with PDB IDs 1JMQ, 1VII, and 2M8I^29,30,31^. For each fold, we generated Potts models with TERMinator and COORDinator and used these to score each mutated protein sequence. We then computed the Pearson correlation between the predicted energies and the experimental stability values.

#### Fine-tuning

For fine-tuning, we froze the weights in all layers except the last output layer. By re-training only the output layer, we avoided both overfitting to the new data and erasing what the model had learned about proteins in the earlier layers. We used the Bcl-2 binding affinity dataset^19^ and associated complex structures to fine-tune the model. We used the Mcl-1 binding data as training data and the Bcl-x_L_ data as a validation, to stop training at the appropriate time. As the loss function, we used a mixed objective between the composite psuedolikelihood and the Pearson correlation between the experimentally measured binding affinities and predicted Potts model energies, weighted 1:100 as discussed in the Loss Functions portion of Section 4.3.

## Acknowledgements

This work was funded by an award from the National Institutes of General Medical Sciences to Gevorg Grigoryan and Amy E. Keating, 5R01GM132117. The authors acknowledge Dartmouth Anthill, MGHPCC C3DDB, and MIT SuperCloud for providing high-performance computing resources used to generate research results for this paper. Alex J. Li acknowledges funding from MIT UROP. Alex J. Li and Mindren Lu would like to thank Sebastian Swanson for teaching them how to use the dTERMen software suite. Vikram Sundar acknowledges funding from the Fannie and John Hertz Foundation. Gevorg Grigoryan is a co-founder, share holder, and Chief Technology Officer at Generate Biomedicines, Inc.

## A Potts Model

Our model outputs a Potts model over positional amino acid labels, commonly known as an energy table. A Potts model describes a mapping from sequence *S* of length *L* to energy *E*(*S*) with the functional form

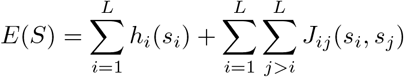

where

- Singleton terms *h_i_*(*s_i_*) describe the energy contribution of position *i* in *S*.
- Pairwise interaction terms *J_ij_*(*s_i_*, *s_j_*) describe the energy contribution from the interaction between positions *i* and *j* in *S*.

In our Potts model, the singleton term takes the form *h_i_*(*s_i_*) = *E_s_*(*R_i_* = *m*), where *E_s_* is a lookup table of energies for placing residue *m* at position *i*. The pairwise interaction term takes the form *J_ij_*(*s_i_*, *s_j_*) = *E_p_*(*R_i_* = *m*, *R_j_* = *n*), where *E_p_* is a lookup table of energies of placing residue *m* at position *i* and residue *n* at position j. This functional form is attractive because it can be used to rapidly evaluate the energy of any sequence, and it is easy to optimize via MCMC-based methods.

## B TERM data

Our goal was to feed into our neural network similar data as would normally be mined as part of dTERMen^9^. This would enable us to differentiate the limitations of TERM data themselves from the limitations associated with the specific statistical approach in dTERMen. To this end, we modified an in-house version of the dTERMen program with the ability to output TERM match information for all of the motifs used in the standard procedure (as described in Zhou et al.^9^). Briefly, dTERMen defines three types of TERMs in the input structural template: singleton, near-backbone, and pair TERMs. Singleton TERMs are defined around each residue *i* via the contiguous fragment between residues (*i* – *n*) and (*i* + *n*), where *n* is a parameter (*n* =1 was used in this study). Near-backbone TERMs combine the local backbone around residue *i* (i.e., the singleton fragment) with local backbone fragments around each residue *j* whose backbone is geometrically poised to interfere with amino-acid sidechains at _i_. Finally, pair TERMs are defined around each pair of residues *i* and *j* that are geometrically positioned to affect each other’s amino-acid choice.

As described in Zhou et al. ^9^, it is frequently the case that the full near-backbone TERM around a residue (i.e., the generally multi-segment motif that captures all relevant surrounding backbone fragments) does not contain sufficient structural matches in the database to generate reliable statistics, in which case the dTERMen procedure seeks to optimally partition the overall near-backbone contribution into as few sub-motifs as possible. This step adds considerable search time. We reasoned that a learning-based approach may be better at extracting relevant statistical couplings between residue sites, such that a detailed breakdown of sidechain-to-sidechain versus backbone-to-sidechain coupling statistics may not be necessary. Therefore, we omitted near-backbone TERMs in this study for computational efficiency.

In finding close structural matches, dTERMen uses a motif complexity-based empirical RMSD cutoff (defined in Mackenzie et al.^10^), with additional settings used to control the minimal and maximal number of matches. In this study, we set these limits to lower values than previously reported^9^ for computational efficiency. Specifically, the minimal/maximal match counts were 200/500 for singleton TERMs, and 400/500 for pair TERMs. Under these settings, dTERMen takes roughly 4 minutes per residue (single-core, 8GB RAM). The native sequence recovery rate of dTERMen was estimated on the basis of energy tables produced with these settings. As input into the neural-network models, only data from the top 50 TERM matches were used.

**FIGURE B1.**
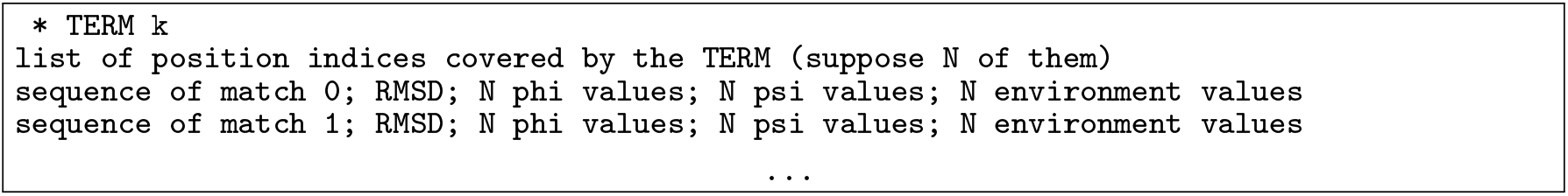
TERM matches information structure.

For each considered TERM, we output which positions of the structural template it covers along with information on each of its matches in the database of known structures, _i_.e. the match sequence, best-fit backbone RMSD from the query, backbone *ϕ* and *ψ* values at each residue, and the “environment” of each residue–a scalar ranging from 0 to 1 that describes how solvent-exposed the residue is (the freedom metric defined in^12^); see Figure B1.

Additionally, for every residue in a TERM we compute a “contact index” that specifies the sequence distance from a central residue in the TERM. TERMs are constructed either around a single central residue (for a singleton TERM) or a pair of residues (for a pair TERM). We assign central residues an index of 0 and define the contact index for the remaining residues as the directional sequence distance to the closest intra-chain center residue. More specifically, non-central residues closer to the N-terminus than their corresponding central residue are assigned a negative integer contact index, while non-central residues closer to the C-terminus than their corresponding central residue are assigned a positive integer contact index.

## C Weighted Cross-Covariance Matrix Features

For each pair of residues, we compute the weighted cross-covariance matrix between the residue features across all matches to the TERM, weighted by RMSD. Let *r_i_* represent the RMSD of match *i* of *n* to the TERM. Then, the weight of match *i* is computed as

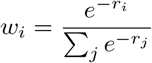

Consider a TERM edge from residue *a* to residue *b*. For match *i*, residue a has *a* vector of features *m_a,i_* and residue *b* has a vector of features *m_b,i_*. This vector includes the one-hot encoding of the residue identity, sinusoidally encoded torsion angles, RMSD, and environment value. The weighted mean of features for residue a and residue *b*, *μ_m_a,i__* and *μ_m_b,i__* are

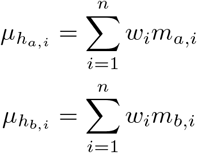

The cross-covariance matrix is then computed as

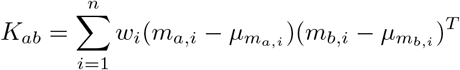

This matrix is then flattened into a vector and its dimensionality is reduced using a two-layer feedforward network with ReLU activations. This output is used as the edge feature between the two residues of concern in the TERM graph.

## D TERM MPNN

We define the following notation:

- *h_i,t_*: the embedding for residue *i* in TERM *t*
- *h_i→j,t_*: the embedding for directional edge *i* → *j* in TERM *t*
- *f_n_, f_e_*: three-layer dense networks with ReLU activations
- *g_n_, g_e_*: two-layer dense networks with ReLU activations
- 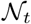 the set of residues in TERM *t*
- *s_i,t_*: the sinusoidal embedding of the contact index of residue *i* in TERM *t*. We multiply the contact index by 500 before performing the sinusoidal embedding, which we find empirically allows for better learning (see positional embedding in Vaswani et al.^26^).
- [;] represents the concatenation operation

The TERM MPNN utilizes alternative edge-update and node-update layers. The update for a directional edge *i → j* in TERM *t* is computed as

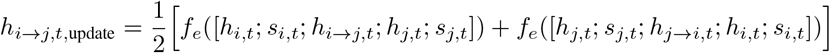

And this update is applied as follows:

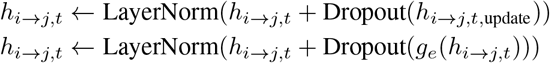

The update for a node is computed as

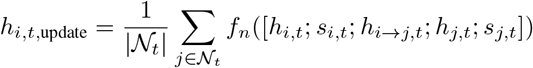

And this update is applied as follows:

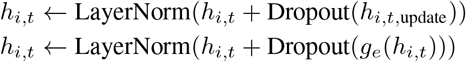

The TERM MPNN contains three layers, with each layer containing an edge update followed by a node update. After these updates, all bidirectional edges are merged into undirected edges via taking the mean of the two edge embeddings.

## E GNN Potts Model Encoder

The GNN Potts Model Encoder is another message-passing network that is identical to the TERM MPNN in architecture but takes in different input features. The GNN Potts Model Encoder operates on a *k*-NN graph rather than a fully-connected graph, meaning node updates are computed over a residue’s *k* nearest neighbors. Additionally, because there is no notion of “contact index” when it comes to global structure, the update function does not take such features as inputs.

Before running message passing, the GNN must stitch together the TERM-based structure embedding and the coordinate-based structure embeddings. Node embeddings for the GNN are computed by concatenating the coordinate-based features from Ingraham et al. ^5^ and the TERM-based features and feeding that vector through a linear layer to compress the vector back to the original dimensionality. Edge embeddings are also formed by computing the coordinate-based edge embedding from Ingraham et al.^5^, concatenating the corresponding TERM edge embeddings, and feeding that vector through a linear layer to compress the vector back to the original dimensionality. In the case that a TERM edge embedding does not exist for that particular *k*-NN graph edge, a zero-vector of equal dimensionality is used instead.

The edge embeddings derived after message-passing is completed are then projected to a 400-dimensional vector by a feedforward network and reshaped to form a matrix containing interaction energies between pairs of interacting residues. The interaction energy matrices give the pair energies of the Potts model. Due to the inclusion of self-edges in the *k*-NN graph, we can also compute self-energies for the Potts model by taking the diagonal of the self-interaction matrix produced by the self-edge for each residue.

**FIGURE E1.**
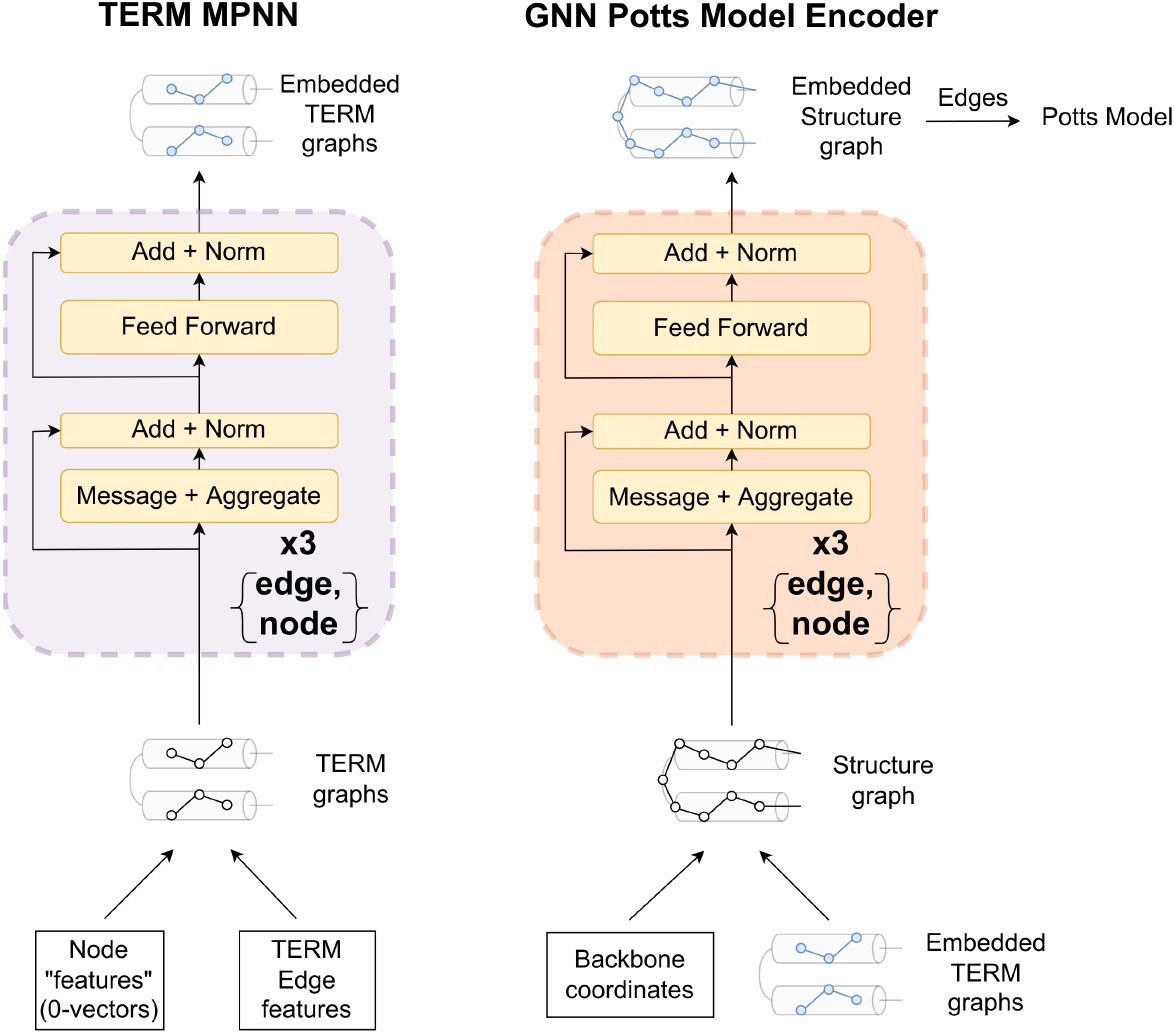
TERMinator Submodule Architectures.

## F Hyperparameters

During training we shuffle according to the semi-shuffle method described in Methods and Materials. We use partition sizes of 500 and perform variable-sized batching, with the cutoff being 55000 TERM residues for TERMinator and 6000 residues for COORDinator.

The GNN Potts Model Encoder has a hidden dimensionality of 128, and consists of three node update layers and three edge update layers which are interleaved. Each of these layers use message-computation layers *f* which are three-layer dense networks with hidden layer sizes (384, 128, 128), as well as feedfoward layers *g* which are two-layer dense networks with hidden layer sizes (512, 128).

The TERM Information Condenser uses a hidden dimension of 32. The cross-covariance matrix uses a 2-layer feedforward network with hidden layer sizes (128, 32). The TERM MPNN consists of three node update layers and three edge update layers which are interleaved. Each of these layers use message-computation layers *f* which are three-layer dense networks with hidden layer sizes (96, 32, 32), as well as feedfoward layers *g* which are two-layer dense networks with hidden layer sizes (128, 32).

The choice of different hidden dimensions for the TERM Information Condenser and the GNN Potts Model Encoder is largely due to GPU memory issues, as raw TERM data are much larger than coordinate data. Given more compute power, one direction to explore is how the model’s performance is affected by varying the hidden dimensionality of both portions of the network.

Across both networks, we use a rate of 0.1 for all Dropout layers.

## G Description of Ablated Models

The following list provides a brief description of TERMinator ablation models:

- **TERM Information Condenser + GNN Potts Model Encoder:** All neural modules included.
- **Ablate TERM Information Condenser:** The TERM Information Condenser is reduced to a series of linear transformations.
  **– Ablate TERM MPNN:** Initial singleton and pairwise TERM features are directly passed to the GNN Potts Model Encoder.
- **Ablate GNN Potts Model Encoder:** Outputs of the TERM Information Condenser are embedded on a *k*-NN graph and then projected to form a Potts Model.
- **Ablate Coordinate-based Features, Retain *k*-NN graph:** Coordinate-based features are set to 0. The *k*-NN graph is still retained.
- **GNN Potts Model Encoder Alone (no TERM information):** Outputs of the TERM Information Condenser are set to 0.

### Ablate GNN Potts Model Encoder vs. dTERMen

In the main text, we claim that the version of TERMinator with the GNN Potts Model Encoder ablated has access to essentially the same features as the version of dTERMen that we call dTERMen*. It is important to acknowledge a few differences in the precise inputs. Regarding TERMinator, while this particular ablation form does not have access to coordinates directly, it does have access to a *k*-NN graph, which dTERMen* does not get. However, when we ablate the GNN Potts Model Encoder, it has no opportunities to perform graph operations over the *k*-NN graph; instead, the *k*-NN graph is only used to restrict pair interactions in the Potts model to those present in the graph. Due to the nature of TERMs being constructed out of small sets of spatially-proximal residues (<7 residues), it is almost always the case that all TERM edges will be included in the *k*-NN graph (in this work, *k* = 30), leading to negligible utilization of the *k*-NN graph itself. On the other hand, dTERMen* used more matches to compute the Potts model than does TERMinator, which was restricted to using the top 50 matches. We also note that the published version of dTERMen uses near-backbone TERMs that were not included for the dTERMen* sequence recovery results reported here, although this also means that TERMinator did not have access to these TERMs either. All things considered, it is reasonable to assume that this ablation of TERMinator and dTERMen* effectively have access to the same types of information, with dTERMen* performing worse despite having access to more matches.

## H Sequence Complexity Plots

**FIGURE H1.**
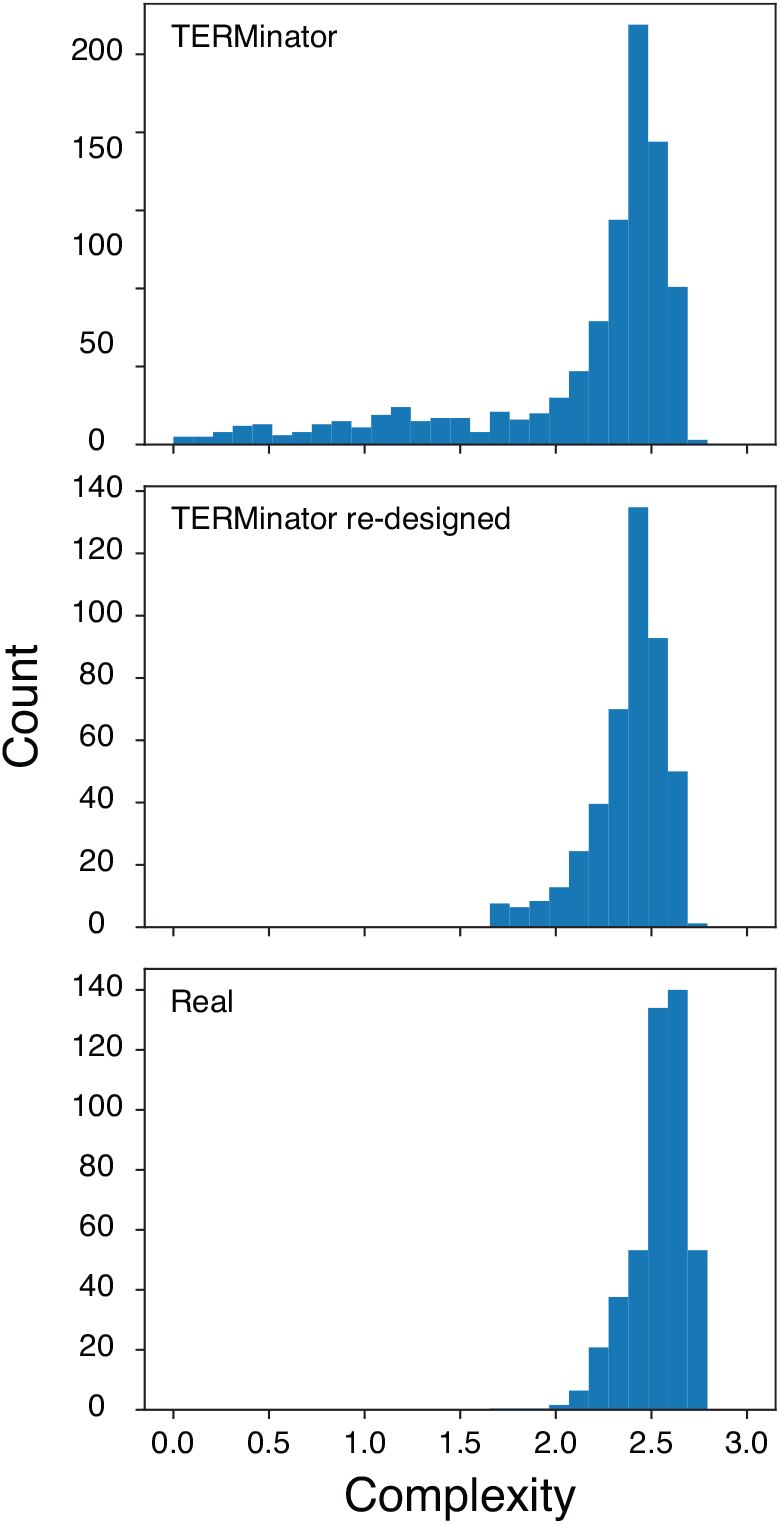
The complexity distribution, based on the number of unique arrangements of labels, of both native and designed sequences for structures in the Ingraham Dataset test set (N=1120). TERMinator re-designed refers to design using the low-complexity penalty, for low-complexity sequences.

**FIGURE H2.**
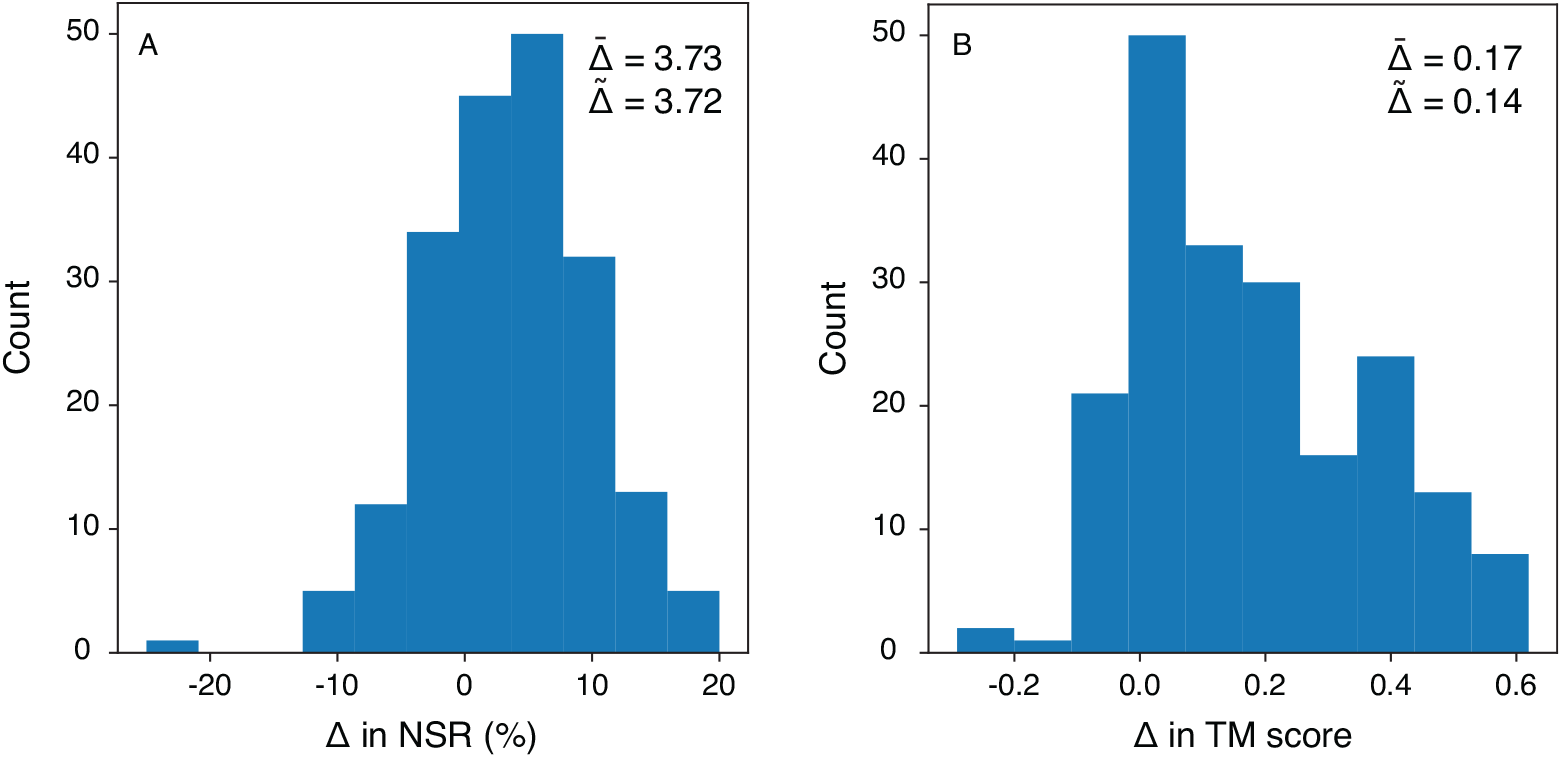
The changes in (A) native sequence recovery (NSR), and (B) TM-score, after re-designing the Ingraham test set low-complexity cases with the complexity-based penalty. 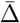 represents the mean change and 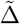 represents the median change, in both graphs, both in percentage points.

**FIGURE H3.**
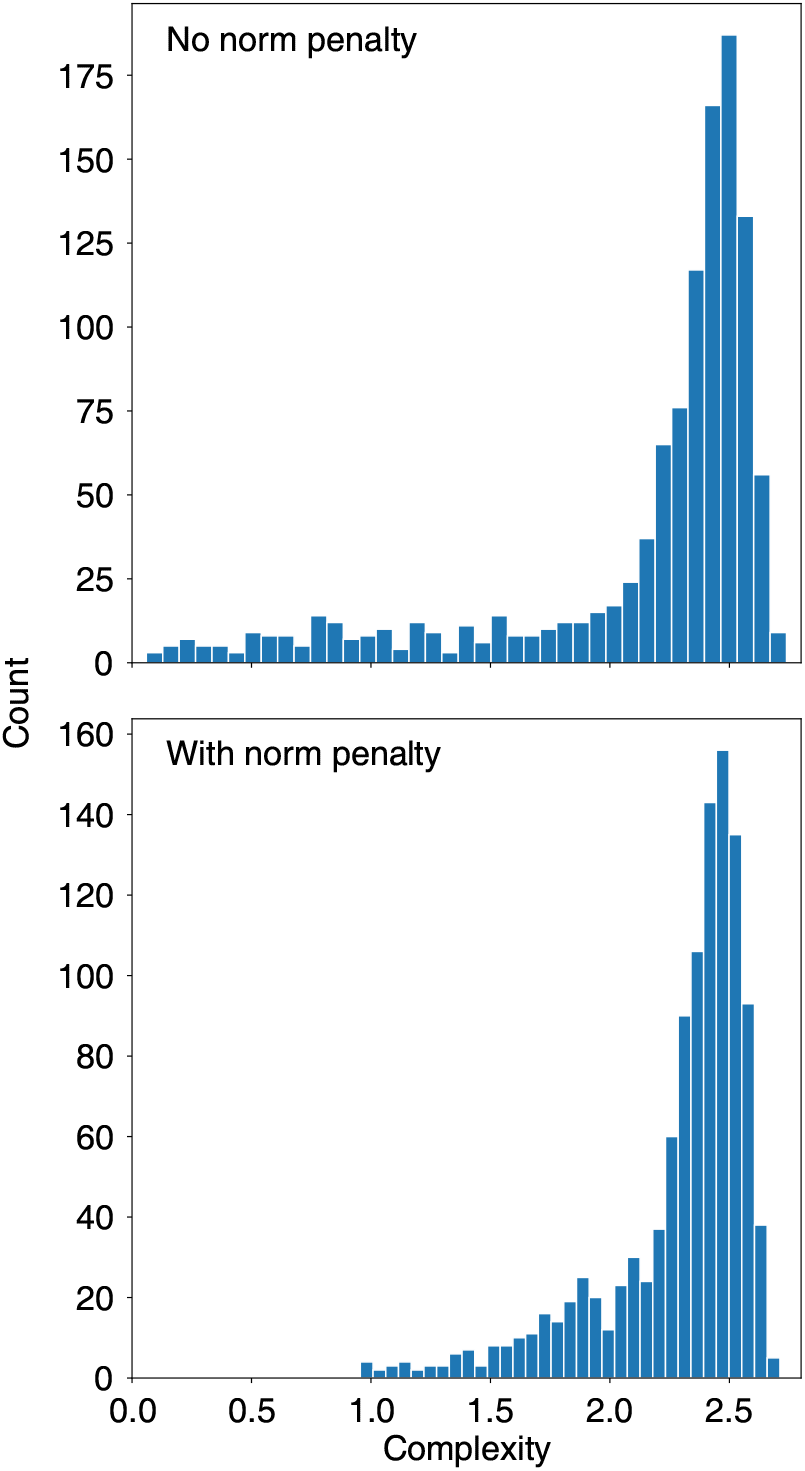
The change in complexity after re-designing the Ingraham test set low-complexity cases with the complexity-based penalty.

## I Folding examples

**FIGURE I1.**
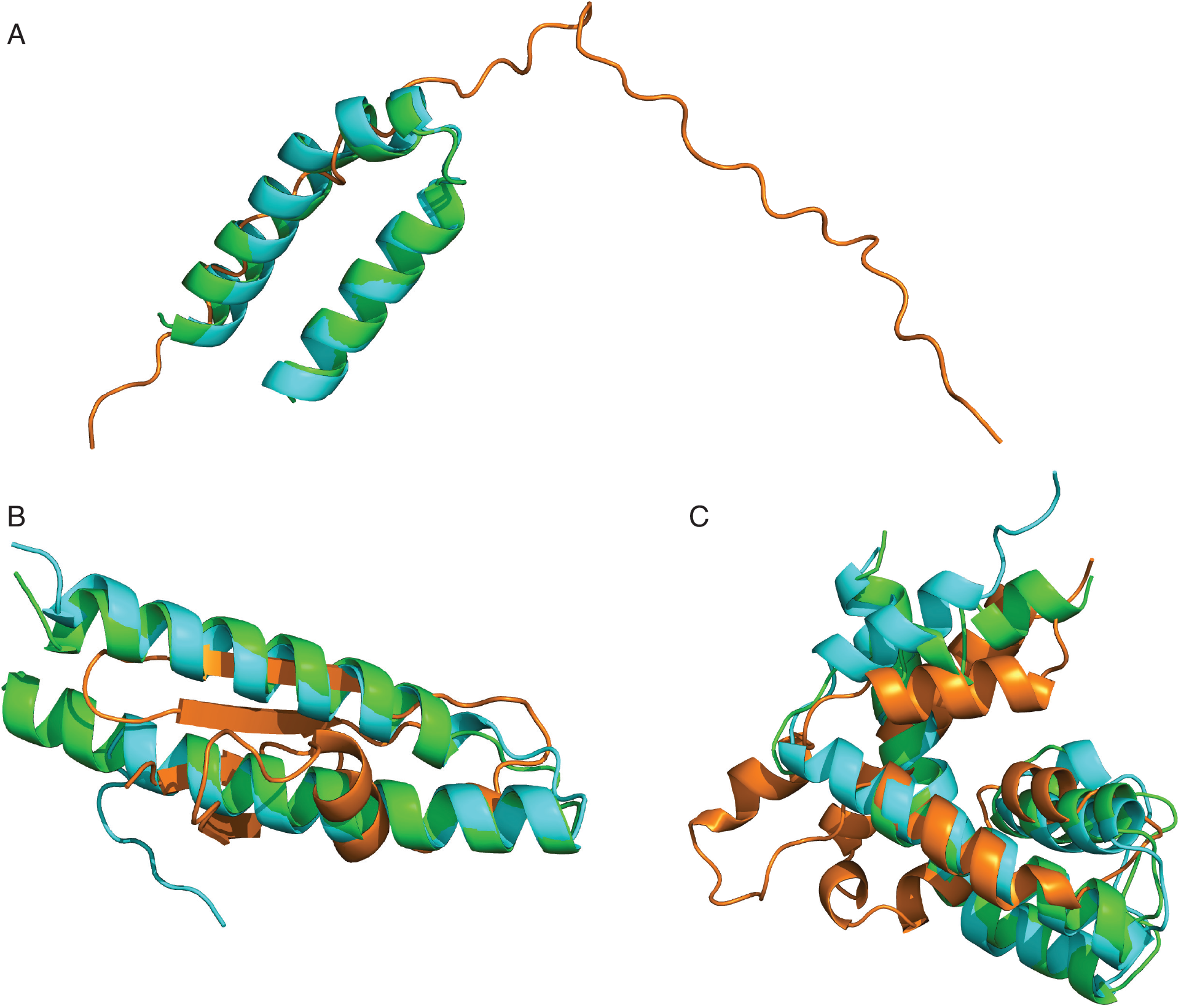
Examples of TERMinator (including low-complexity sequence re-desgin) recovering the correct fold at low NSR compared to randomized sequences at similar NSR. A) 1NVP_B - TERMinator predicted a sequence with NSR=16.28%; the AlphaFold (AF) structure (cyan) had TM-score=0.83 and pLDDT=90.26. The randomized control with NSR=18.6% had a structure (orange) with TM-score=0.2 and pLDDT=72.09, B) 1SKV_A - TERMinator predicted a sequence with NSR=18.75%, and the AF structure (cyan) had TM-score=0.81 and pLDDT=87.73. The control with NSR=18.75% had a structure (orange) with TM-score=0.25 and pLDDT=50.63, C) 5CBG_A - TERMinator predicted a sequence with NSR=18.63%, and AF structure (cyan) had TM-score=0.81 and pLDDT=78.56. The control with NSR=18.63% had a structure (orange) with TM-score=0.4 and pLDDT=60.57.

**FIGURE I2.**
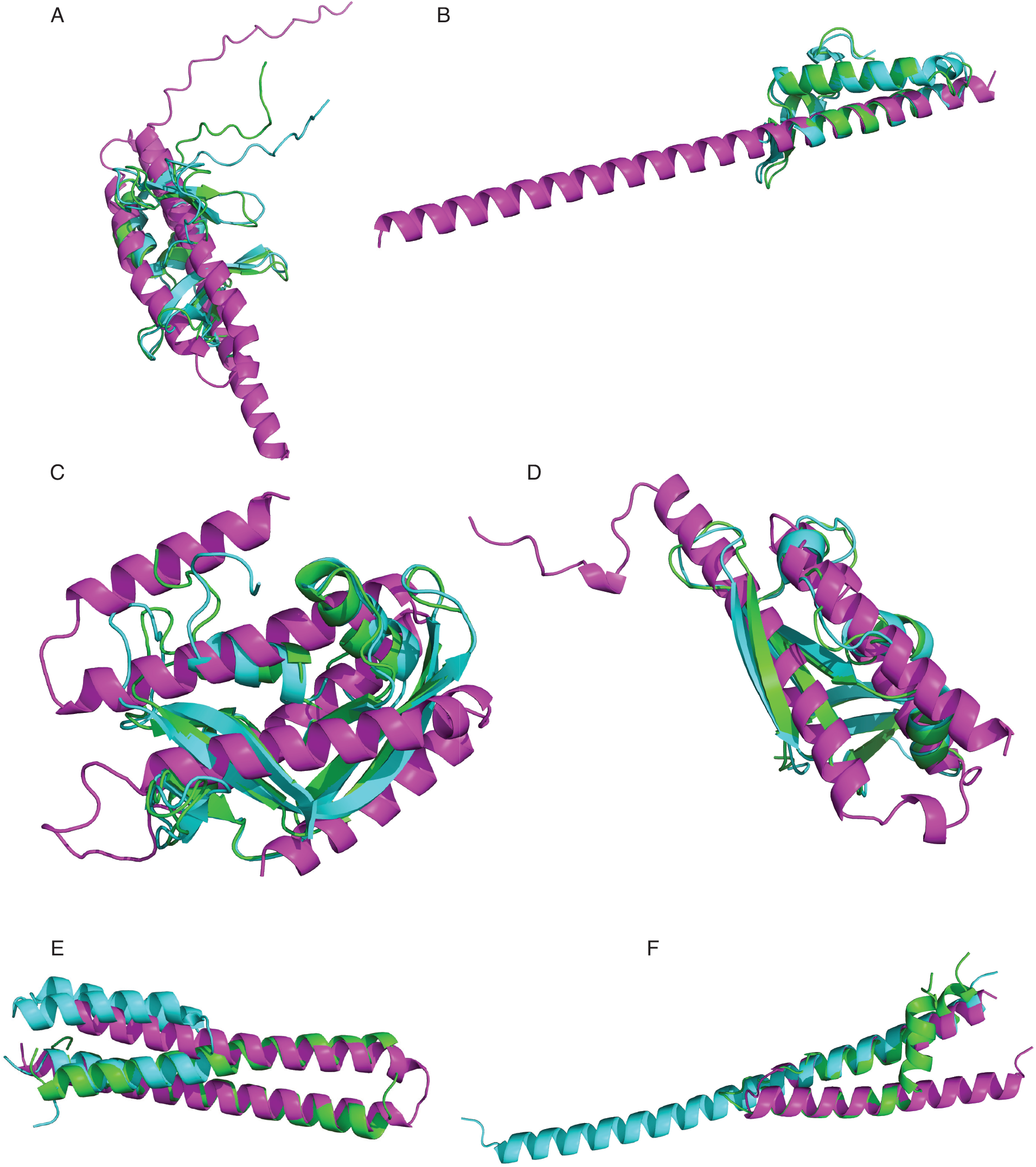
Examples of extreme changes in fold specificity before (magenta) and after (cyan) TERMinator redesign with the complexity penalty are shown here after being aligned (using TMalign) to the native chain (shown in green). A) 1YUA_A had a TM-score of 0.23 and improved to a TM-score of 0.79 (240%); the NSR improved from 27.05% to 31.15%, B) 1SG7_A TM-score improved from 0.30 to 0.87 (187%) and NSR improved from 16.0% to 21.33%, C) 2HKY_A TM-score improved from 0.26 to 0.84 (227%) and NSR improved from 19.38% to 27.91%, D) 1KVZ_A TM-score improved from 0.26 to 0.88 (236% increase) and NSR improved from 20.56% to 28.04%, E) 4I0X_L TM-score dropped from 0.65 to 0.39 (41%) and NSR dropped from 25.61% to 23.17%, and F) 1BCC_H TM-score dropped from 0.63 to 0.34 (−46% change) but NSR improved from 18.18% to 30.30%.

## J Full Results for Energy-Based Benchmarks

Here, we report the individual values for the performance of TERMinator and COORDinator on the energy-based protein analysis tasks discussed in the Results section of this paper.

**TABLE J1.** Affinity correlation performance averaged over all Bcl-2 complex templates^13,19^. These are the summary values used to generate Figure 3.

|  | **Bcl-x<sub>L</sub>** | **Bfl-1** | **Mcl-1** | **Mean** |
| --- | --- | --- | --- | --- |
| FoldX | 0.23 ± 0.11 | 0.34 ± 0.07 | 0.37 ± 0.11 | 0.31 ± 0.06 |
| Rosetta | 0.24 ± 0.06 | 0.32 ± 0.03 | 0.45 ± 0.04 | 0.34 ± 0.09 |
| dTERMen | 0.27 ± 0.06 | 0.41 ± 0.01 | 0.45 ± 0.08 | 0.38 ± 0.08 |
| TERMinator, Single-Chain | 0.13 ± 0.10 | 0.32 ± 0.07 | 0.47 ± 0.11 | 0.30 ± 0.14 |
| TERMinator, Multi-Chain | 0.22 ± 0.12 | 0.36 ± 0.08 | 0.46 ± 0.13 | 0.35 ± 0.10 |
| COORDinator, Single-Chain | 0.12 ± 0.11 | 0.27 ± 0.08 | 0.41 ± 0.09 | 0.26 ± 0.12 |
| COORDinator, Multi-Chain | 0.18 ± 0.10 | 0.36 ± 0.09 | 0.45 ± 0.11 | 0.33 ± 0.11 |

**TABLE J2.** Protein stability correlation performance on a mutational stability dataset of *de novo* small proteins^21^, including previous work^5 22^. For all proteins and both TERMinator and COORDinator, the standard deviations across triplicate training runs were ≤ 0.04 and are omitted for clarity. The top half of the table contains the values used to generate Figure 4. The bottom half of the table notes our performance on 7 additional structures in the Rocklin et al. ^21^ dataset for which the other models do not report results.

|  | **Structured**<br>**GNN<sup>5</sup>** | **GVP-**<br>**GNN<sup>6</sup>** | **GVP+**<br>**AF2<sup>22</sup>** | **Single-Chain**<br>**TERMinator** | **Multi-Chain**<br>**TERMinator** | **Single-Chain**<br>**COORDinator** | **Multi-Chain**<br>**COORDinator** |
| --- | --- | --- | --- | --- | --- | --- | --- |
| $\beta\beta\alpha\beta\beta_{37}$ | 0.50 | 0.53 | 0.70 | 0.57 | 0.55 | 0.60 | 0.57 |
| $\beta\beta\alpha\beta\beta_{1498}$ | 0.44 | 0.39 | 0.33 | 0.38 | 0.34 | 0.37 | 0.34 |
| $\beta\beta\alpha\beta\beta_{1702}$ | 0.17 | 0.26 | 0.22 | 0.26 | 0.21 | 0.23 | 0.19 |
| $\beta\beta\alpha\beta\beta_{1716}$ | 0.40 | 0.57 | 0.58 | 0.54 | 0.51 | 0.56 | 0.48 |
| $\alpha\beta\beta\alpha_{779}$ | 0.56 | 0.48 | 0.64 | 0.58 | 0.51 | 0.58 | 0.56 |
| $\alpha\beta\beta\alpha_{223}$ | 0.33 | 0.47 | 0.55 | 0.47 | 0.44 | 0.51 | 0.46 |
| $\alpha\beta\beta\alpha_{726}$ | 0.21 | 0.19 | 0.26 | 0.22 | 0.21 | 0.23 | 0.19 |
| $\alpha\beta\beta\alpha_{872}$ | 0.23 | 0.39 | 0.42 | 0.33 | 0.27 | 0.36 | 0.29 |
| $\alpha\alpha\alpha_{134}$ | 0.36 | 0.44 | 0.50 | 0.45 | 0.44 | 0.46 | 0.43 |
| $\alpha\alpha\alpha_{138}$ | 0.41 | 0.44 | 0.58 | 0.48 | 0.45 | 0.49 | 0.45 |
| **Mean** | 0.36 | 0.42 | 0.48 | 0.43 | 0.39 | 0.44 | 0.40 |
| $\alpha\alpha\alpha_{142}$ | | | | 0.65 | 0.61 | 0.65 | 0.62 |
| $\beta\beta\alpha\beta_{882}$ | | | | 0.61 | 0.56 | 0.60 | 0.55 |
| $\beta\beta\alpha\beta_5$ | | | | 0.43 | 0.44 | 0.45 | 0.45 |
| $\beta\beta\alpha\beta_{15}$ | | | | 0.38 | 0.36 | 0.37 | 0.36 |
| 1JMQ |  |  |  | 0.45 | 0.39 | 0.41 | 0.36 |
| 1VII |  |  |  | 0.33 | 0.29 | 0.40 | 0.22 |
| 2M8I |  |  |  | 0.56 | 0.52 | 0.56 | 0.54 |
| **Mean (all folds)** |  |  |  | 0.45 | 0.42 | 0.46 | 0.42 |

**TABLE J3.** Protein stability correlation performance on a mutational stability dataset of *de novo* small proteins^21^, for the GVP-Transformer-AF2 model from Hsu et al.^22^ and the single-chain TERMinator and COORDinator models, both before and after fine-tuning. The top half of the table includes the 10 structures studied in previous work, and the bottom half of the table notes our performance on 7 additional structures from Rocklin et al. ^21^ which have not been previously benchmarked.

|  | **GVP+AF2<sup>22</sup>** | **Single-Chain**<br>**TERM** | **TERMinator**<br>**Fine-Tuned** | **Single-Chain**<br>**COORD** | **COORDinator**<br>**Fine-Tuned** |
| --- | --- | --- | --- | --- | --- |
| $\beta\beta\alpha\beta\beta_{37}$ | 0.70 | 0.57 | 0.64 | 0.60 | 0.66 |
| $\beta\beta\alpha\beta\beta_{1498}$ | 0.33 | 0.38 | 0.50 | 0.37 | 0.50 |
| $\beta\beta\alpha\beta\beta_{1702}$ | 0.22 | 0.26 | 0.29 | 0.23 | 0.27 |
| $\beta\beta\alpha\beta\beta_{1716}$ | 0.58 | 0.54 | 0.63 | 0.56 | 0.64 |
| $\alpha\beta\beta\alpha_{779}$ | 0.64 | 0.58 | 0.68 | 0.58 | 0.69 |
| $\alpha\beta\beta\alpha_{223}$ | 0.55 | 0.47 | 0.59 | 0.51 | 0.60 |
| $\alpha\beta\beta\alpha_{726}$ | 0.26 | 0.22 | 0.43 | 0.23 | 0.44 |
| $\alpha\beta\beta\alpha_{872}$ | 0.42 | 0.33 | 0.53 | 0.36 | 0.56 |
| $\alpha\alpha\alpha_{134}$ | 0.50 | 0.45 | 0.46 | 0.46 | 0.50 |
| $\alpha\alpha\alpha_{138}$ | 0.58 | 0.48 | 0.50 | 0.49 | 0.52 |
| **Mean** | **0.48** | **0.43** | **0.53** | **0.44** | **0.54** |
| $\alpha\alpha\alpha_{142}$ | | 0.65 | 0.69 | 0.65 | 0.71 |
| $\beta\beta\alpha\beta_{882}$ | | 0.61 | 0.62 | 0.60 | 0.63 |
| $\beta\beta\alpha\beta_5$ | | 0.43 | 0.51 | 0.45 | 0.53 |
| $\beta\beta\alpha\beta_{15}$ | | 0.38 | 0.41 | 0.37 | 0.40 |
| 1JMQ |  | 0.45 | 0.49 | 0.41 | 0.46 |
| 1VII |  | 0.33 | 0.47 | 0.40 | 0.50 |
| 2M8I |  | 0.56 | 0.60 | 0.56 | 0.62 |
| **Mean (all folds)** |  | **0.45** | **0.53** | **0.46** | **0.54** |

## K Potts Model Parameter Distributions for TERMinator and COORDinator

**FIGURE K1.**
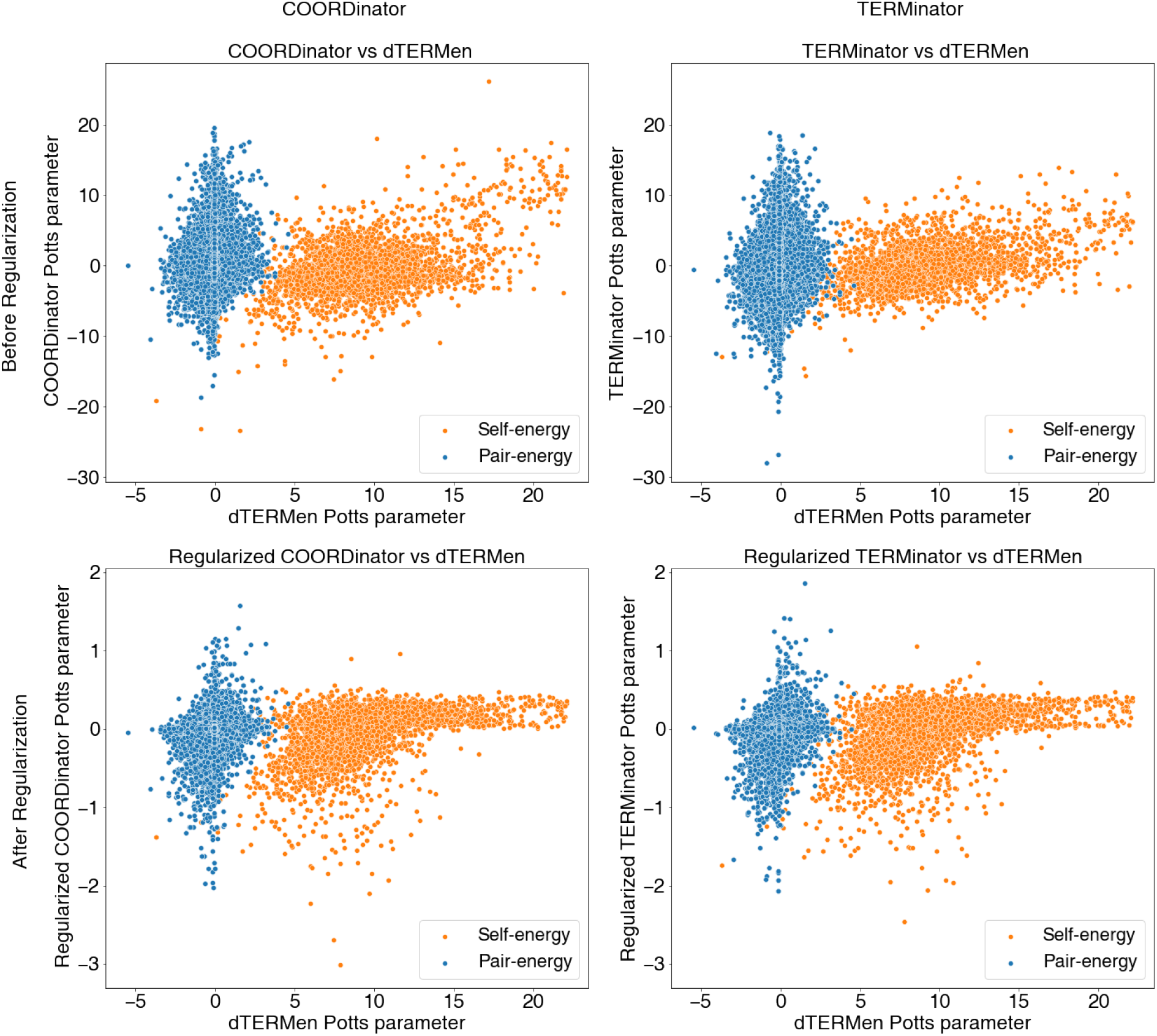
TERMinator and COORDinator Potts model parameters versus dTERMen* Potts model parameters for a representative test case, before and after regularization.

